# Acinar-Derived VEGF-A Orchestrates Blood Vascular Remodeling and Preserves Microvessels During Acute Pancreatitis

**DOI:** 10.1101/2025.07.06.663280

**Authors:** Elias Aajja, Hélène Lefort, Siam Mahibullah, Doriane Frederic, Laura Vandooren, Patrick Henriet, Donatienne Tyteca, Christophe E. Pierreux

## Abstract

Acute pancreatitis is a common inflammatory condition of the pancreas that can lead to severe complications such as chronic pancreatitis and pancreatic ductal adenocarcinoma. While vascular remodeling is a hallmark of many inflammatory conditions, the molecular mechanisms underlying vascular changes during pancreatitis remain largely unexplored. This study aimed to investigate the vascular changes associated with acute pancreatitis and to identify the molecular mechanisms underlying these changes.

Acute pancreatitis was induced in wild-type mice through caerulein injections and resulted in progressive and substantial vascular changes, characterized by morphological remodeling, increased vessel density, and elevated vascular permeability. These structural changes were accompanied by molecular alterations, including increased expression of endothelial genes and decreased surface expression of vascular endothelial VE-cadherin. Injured acinar cells exhibited a significant increase in VEGF-A expression during pancreatitis. Acinar-specific VEGF-A inactivation led to marked impairments in vascular remodeling, with reduced vessel density and diminished number of vessels, without affecting immune cell infiltration, fibrosis, or acinar-to-ductal metaplasia.

Together, our work identifies a mechanism by which increased expression of VEGF-A by acinar cells during pancreatitis is essential for maintaining vascular integrity and for driving vascular remodeling in the inflamed pancreas.

## Background & Aims

Acute pancreatitis is a serious and increasingly common gastrointestinal disease characterized by a rapid but transient inflammation of the pancreatic tissue. The Western lifestyle greatly increasing the likelihood of developing this condition and common risk factors include alcohol, gallstones, and hypertriglyceridemia [1–3]. While most patients recover quickly when the insult disappears, some may face serious complications such as chronic pancreatitis, pancreatic insufficiency, diabetes mellitus, or even pancreatic cancer [4]. Indeed, pancreatitis is a well-known risk factor for pancreatic ductal adenocarcinoma (PDAC), the most common and deadliest form of pancreatic cancer. This underscores the importance of understanding how acute inflammation can evolve into chronic disease and potentially initiate oncogenesis.

The initial molecular trigger for acute pancreatitis is the premature activation of digestive enzymes, leading to pancreas autodigestion and acinar cell death. These events initiate an inflammatory reaction that in cascade profoundly remodels the pancreatic parenchyma [1–3]. The three commonly described tissular and morphological changes in acute pancreatitis are immune cell infiltration, collagen deposition leading to fibrosis, and acinar cell dedifferentiation into ductal-like cells also called acinar-to-ductal metaplasia (ADM) [5–8]. This latter may be a protective mechanism to limit further release and premature activation of digestive enzymes [7, 9, 10]. However, persistent ADM under inflammatory conditions can act as a precursor to PDAC, highlighting the fine balance between regeneration and chronicity in this disease [11].

During inflammation, blood vessels are central actors that undergo significant changes, such as angiogenesis, vascular remodeling, increased permeability, and vasodilation [12–14]. Historically, vascular changes during pancreatitis were described in the 1980s and 1990s through experimental models, but these studies largely focused on the morphological level, without delving into the molecular mechanisms involved [15–18]. These vascular changes are well-documented in other inflammatory diseases, such as inflammatory bowel disease, where Vascular Endothelial Growth Factor A (VEGF-A) plays a pivotal role [19]. VEGF-A is a critical regulator of vascular dynamics during inflammation in other tissues, such as the colon and the lung, as it promotes angiogenesis, increases vascular permeability, and maintains endothelial cell survival via its interaction with VEGF receptor 2 (VEGFR2) [19–21]. VEGF-A is indeed the quintessential angiogenic factor. It not only stimulates angiogenesis, i.e. the formation of new blood vessels, but also maintains homeostasis and/or survival of existing vessels by preventing their regression, a role demonstrated in contexts such as tumor vasculature and retinal angiogenesis [22, 23]. VEGF-A is also known as Vascular Permeability Factor (VPF) and was shown to increase permeability during inflammation in tissues such as the colon and the lung, a process in which VEGF-A is essential for enabling immune cell infiltration and the inflammatory response [19–21].

Blood vascular changes occur during acute pancreatitis, but the cellular and molecular events are poorly described. Moreover, the mechanisms governing these changes remain poorly understood and the role of VEGF-A in acute pancreatitis has not yet been explored. This is particularly relevant since pancreatitis-favored PDAC is characterized by hypovascularization due to vascular collapse, which impairs blood flow and complicates treatment [24]. To fill these gaps in our knowledge, we used a mouse model of acute pancreatitis to describe and quantify the vascular changes during pancreatitis. We then investigated the role of VEGF-A in this model using acinar-specific *Vegfa* knockout mice to uncover the mechanisms through which VEGF-A may contribute to vascular remodeling during acute pancreatitis. We show that blood vascular changes accompany the development of immune cell infiltration, the development of fibrosis, and ADM during acute pancreatitis. We also show that secretion of VEGF-A, by injured acinar cells, controls and maintains blood vessels during pancreatitis.

## Results

### Caerulein-induced acute pancreatitis

Acute pancreatitis was induced in C57BL/6N mice through repetitive caerulein injections, following the treatment timeline shown in **Figure 1A**. Pancreatic tissues were harvested for histological and molecular analyses one day after the final caerulein injection (Day 6; **Fig. 1A**), but also on alternate days after earlier injections (Day 2 and Day 4; **Supplementary Fig. 1A**).

**Figure 1.**
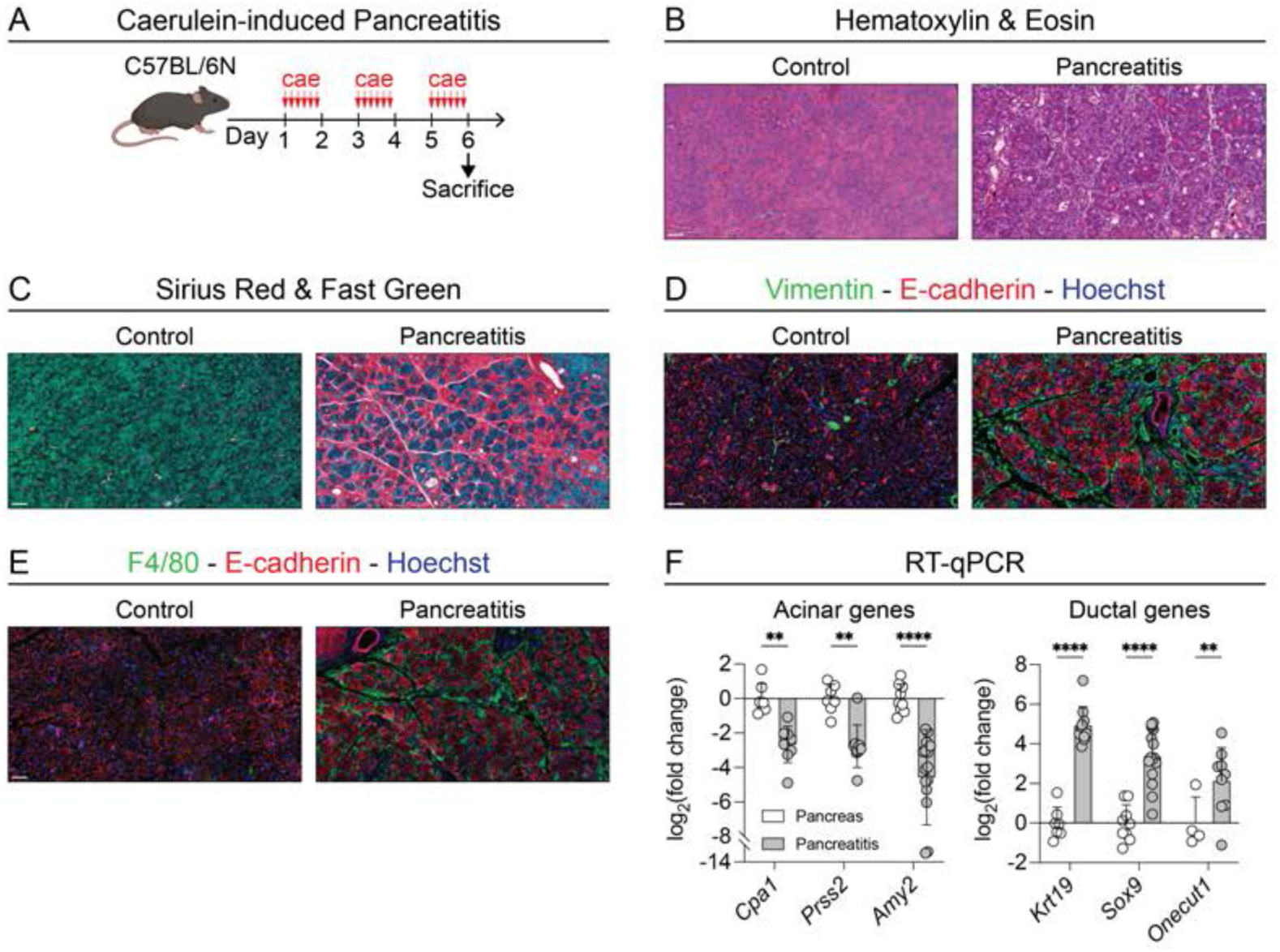
Caerulein-induced model of acute pancreatitis. (A) Timeline of caerulein injections and sacrifice for tissue harvest on day 6 in C57BL/6N mice. Vertical red arrows illustrate the six hourly injections. (B-C) Histological staining of pancreatic sections from control and caerulein-treated (acute pancreatitis) mice: hematoxylin (nuclei) & eosin (cytoplasm) (B), and sirius red (collagen fibers) & fast green (cytoplasm) (C). Scale bar: 50 µm. (D-E) Immunofluorescence localization of vimentin and E-cadherin (for fibroblasts and epithelial cells; D), and of F4/80 and E-cadherin (macrophages and epithelial cells; E) with a nuclear counterstain (Hoechst) in pancreatic sections from control and caerulein-treated mice. Scale bar: 50 µm. (F) Gene expression analysis by RT-qPCR of acinar (*Cpa1*, *Prss2*, *Amy2*) and ductal (*Krt19*, *Sox9*, *Onecut1*) markers, normalized to *18S* ribosomal RNA in pancreatic tissue from control and caerulein-treated mice. Data are shown as means ± SD. Statistical analysis: two-way ANOVA with Holm-Šídák correction. \**p* < 0.05, \*\**p* < 0.01, \*\*\**p* < 0.001, \*\*\*\**p* < 0.0001. n≥4.

The tissular characteristics of pancreatitis developed progressively with the injections. Hematoxylin (nuclei) & eosin (cytoplasm) staining revealed significant parenchymal remodeling, which began at Day 4 and peaked at Day 6, as per our acute pancreatitis protocol (**Supplementary Fig. 1B**). These changes were particularly evident when comparing untreated controls with Day 6 pancreatitis samples (**Fig. 1B**). In controls, the pancreatic parenchyma, particularly the acinar compartment, appeared densely packed and predominantly pink due to abundant apical eosinophilic zymogen granules (**Fig. 1B**). In contrast, tissues affected by pancreatitis exhibited reduced eosinophilia, and increased basophilia (**Fig. 1B** and **Supplementary Fig. 1B**). These changes are consistent with caerulein-induced acinar injury (tissue shrinkage), acinar-to-ductal metaplasia (ADM; reduced eosinophilia) and increased interstitial nuclear density due to immune cell infiltration and fibrosis (increased basophilia; **Fig. 1B**).

Excess deposition of collagen fibers in pancreatitis, visualized using Sirius Red staining, confirmed massive fibrosis at Day 6 (**Fig. 1C**). Fibrosis was further supported by the increased immunofluorescence labeling for vimentin, expressed by fibroblasts, during pancreatitis (**Fig. 1D** and **Supplementary. Fig. 1C**). Immunofluorescence labeling of F4/80 highlighted an increase in macrophage abundance during establishment of acute pancreatitis, confirming inflammation (**Fig. 1E** and **Supplementary Fig. 1D**). Colabeling of the epithelial compartment, using E-cadherin, revealed a remodeled and sparser epithelial compartment in pancreatitis, as compared to controls, confirming epithelial remodeling, likely linked to acinar-to-ductal metaplasia (**Fig. 1D-E** and **Supplementary Fig. 1C-D**). ADM was further investigated by RT-qPCR and this molecular analysis confirmed a decrease in acinar-specific gene expression and an increase in ductal-like gene expression was visible as of Day 4 and more evident at Day 6 (**Fig. 1F** and **Supplementary Fig. 1E**).

In summary, these results indicate that our caerulein-induced pancreatitis model effectively recapitulates the known pathological features of acute pancreatitis, namely immune cell infiltration, fibrosis, and acinar-to-ductal metaplasia.

### Acute pancreatitis is accompanied with blood vascular remodeling

We next investigated the vascular compartment by performing immunofluorescence labeling of endomucin, an endothelial cell marker (**Fig. 2A** and **Supplementary Fig. 2A**). This analysis revealed distinct qualitative morphological changes in blood vessels during pancreatitis as compared to controls (**Fig. 2A**). Notably, these vascular changes progressed over the course of caerulein treatment and were evident as of Day 4 (**Supplementary Fig. 2A**). To quantitatively assess these vascular modifications, the endothelial endomucin-positive objects were segmented (**Fig. 2B** and **Supplementary Fig. 2B**), and morphologically characterized using their area, circularity (**Fig. 2C** and **Supplementary Fig. 2C**), as well as their aspect ratio, and roundness (**Supplementary Fig. 2C**). During pancreatitis, the distribution of the number of endomucin-labeled objects as a function of the area shifted, revealing increased number of small objects (< 80 µm^2^) and conversely, fewer larger ones (> 80 µm^2^). Concomitantly, the circularity (4π × area/perimeter^2^) of these endothelial structures shifted toward values closer to 1, indicating a higher prevalence of more circular structures (**Fig. 2C**). The shift in circularity was apparent by Day 2, while the shift in area became evident by Day 4 (**Supplementary Fig. 2C**). In parallel, the roundness (4 × area / (π × major axis^2^)), increased towards values closer to 1, reflecting a trend towards rounder shapes, while the aspect ratio (major/minor axis), approached 1, indicating a shift towards more equidimensional structures (**Supplementary Fig. 2C**). When plotting circularity as a function of area, with bins color-coded to represent the number of objects, we observed that the increase in small structures and circular structures during pancreatitis corresponded to the same population of objects (**Fig. 2D**; upper left quadrant). Specifically, the rise in smaller structures was driven by an increase in highly circular objects, while the reduction in larger structures primarily involved non-circular objects (**Fig. 2D**; lower right quadrant). This indicates that pancreatitis selectively favors the formation of small, circular structures, while reducing the prevalence of large, non-circular objects.

**Figure 2.**
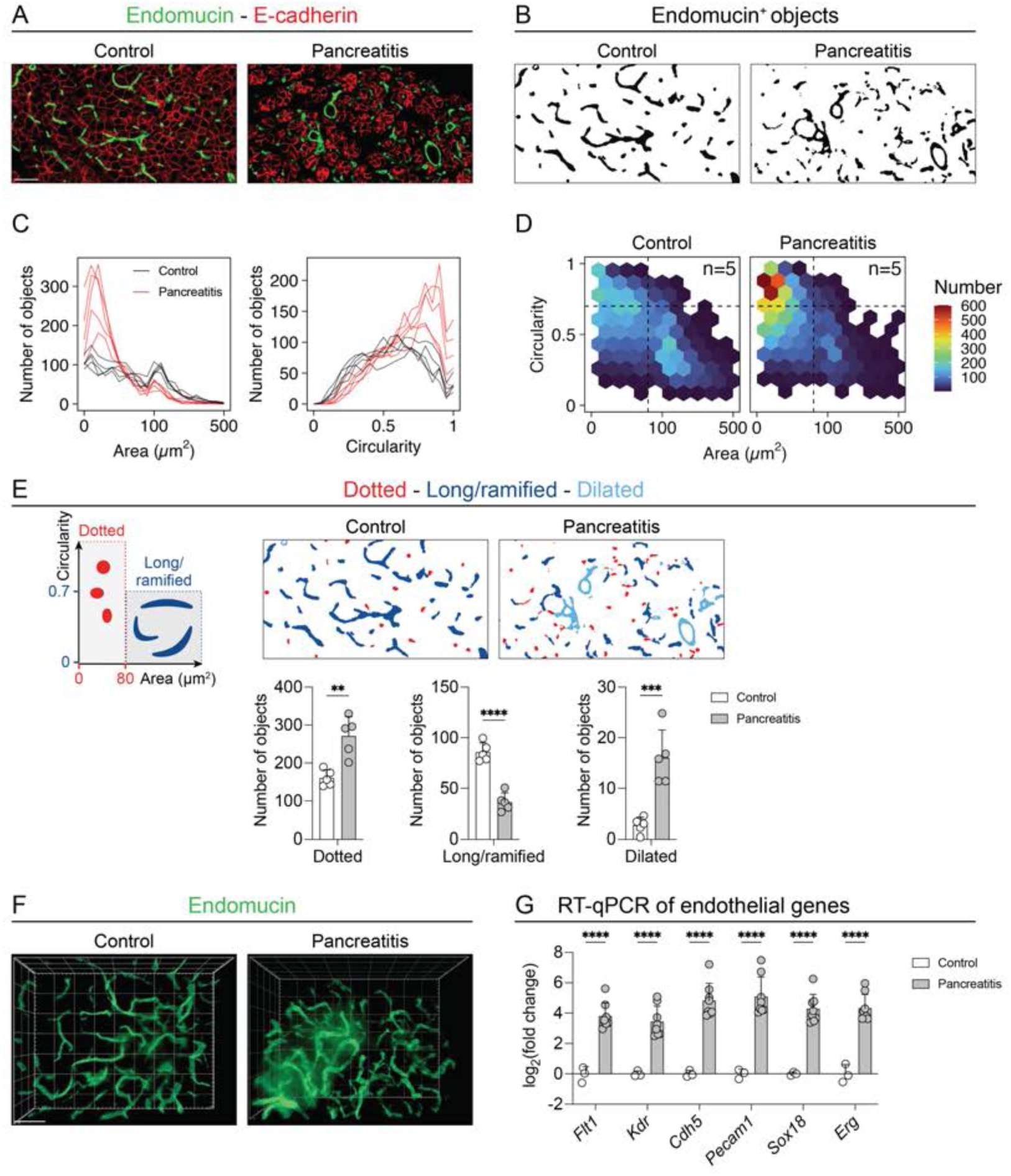
Blood vascular remodeling during acute pancreatitis. (A) Immunofluorescence labeling of endomucin (endothelium) and E-cadherin (epithelial cells) in pancreatic sections from control and caerulein-treated mice. Scale bar: 50 µm. (B) Segmentation of endomucin-positive objects from panel A for morphological analysis. (C) Quantification of endomucin-positive objects: area-based (at left) and circularity-based (at right) 1D count plot in pancreatic sections from control (black curves) and caerulein-treated mice (red curves). Data represent the cumulative counts from five representative 488,160 µm^2^ fields per mouse. n=5. (D) Hexagonal binned plot of circularity vs. area (µm^2^) of endomucin-positive objects, with colors representing object count per bin in pancreatic sections from control and caerulein-treated mice. Data represent the cumulative counts from five representative 488,160 µm^2^ fields per mouse. n=5. (E) Classification of endomucin-positive objects: dotted (area ≤ 80 µm^2^), long/ramified (area > 80 µm^2^, circularity < 0.7), and dilated (manually classified) in pancreatic sections from control and caerulein-treated mice. Data represent the average counts from five representative 488,160 µm^2^ fields per mouse and are shown as means ± SD. Statistical analysis: two-tailed Student’s t-test. \**p* < 0.05, \*\**p* < 0.01, \*\*\**p* < 0.001, \*\*\*\**p* < 0.0001. n=5. (F) 3D reconstruction of vascular structures labeled with endomucin in pancreatic tissue from control and caerulein-treated mice. Scale bar: 50 µm. (G) Gene expression analysis by RT-qPCR of endothelial markers (*Flt1*, *Kdr*, *Cdh5*, *Pecam1*, *Sox18*, *Erg*), normalized to *18S* ribosomal RNA, in pancreatic tissue from control and caerulein-treated mice. Data are shown as means ± SD. Statistical analysis: two-way ANOVA with Holm-Šídák correction. \**p* < 0.05, \*\**p* < 0.01, \*\*\**p* < 0.001, \*\*\*\**p* < 0.0001. n=3 for control, n=8 for pancreatitis.

To further refine the analysis, we classified blood vessels into distinct groups based on their morphology. The less abundant and more complex dilated structures were manually isolated from images and counted separately. Non-dilated objects with an area lower than 80 µm^2^ were categorized as dotted objects, while those with an area larger than 80 µm^2^ and a circularity value below 0.7 were defined as long/ramified objects. Based on this classification, we quantified a significant increase in both dotted and dilated structures, accompanied by a reduction in long/ramified objects (**Fig. 2E**). These changes were observable as of Day 4 (**Supplementary Fig. 2D**). These vascular changes persisted, including the increase in dotted objects and the reduction in long/ramified objects, even when sections were analyzed at high magnifications (**Supplementary Fig. 2E**). This indicates that endothelial cells undergo shape alterations during pancreatitis.

Traditional 2D analyses of tissue provide useful cross-sectional views but they can miss the overall organization of the vascular network. We therefore performed whole-mount 3D imaging, which offers a more comprehensive and volumetric view of the tissue. 3D analysis of pancreatic tissue fragments revealed an increase in vessel density and thickness in pancreatitis, as compared to controls (**Fig. 2F**).

Moreover, RT-qPCR analysis of the endothelial markers *Flt1*, *Kdr, Cdh5*, *Pecam1*, *Sox18*, and *Erg*, revealed striking and consistent (∼16X) increased expression of each of these endothelial genes (**Fig. 2G**). Together, these findings demonstrate that acute pancreatitis induces significant vascular remodeling, characterized by shape alterations, vessel dilation, increased vessel density, and thickness, consistent and concomitant with endothelial cell activation.

### Acute pancreatitis displays dysfunctional blood vessels

Following the morphological and molecular analyses, we next sought to functionally characterize the blood vasculature during pancreatitis. To assess vascular perfusion and permeability, FITC-albumin, a fluorescent tracer, was injected intravenously into mice 10 minutes before sacrifice and analysis. The distribution of FITC-albumin was tracked with fluorescence microscopy. Within this brief 10-minute time-course, FITC-albumin exhibited widespread distribution throughout the entire pancreatic parenchyma, closely mirroring the vascular architecture (compare **Fig. 3A** at left with **Fig. 2A**). In stark contrast, a larger area of the pancreatic parenchyma was occupied by FITC-albumin in caerulein-treated pancreas, suggesting vascular leakage and disrupted vessel integrity upon pancreatitis (**Fig. 3A**, at right). Quantification of the FITC-albumin surface indeed revealed a 6-fold increase when compared to control pancreas (**Fig. 3A**). Colabeling with endomucin and E-cadherin confirmed FITC-albumin confinement within endomucin-positive structures in control pancreas and extravasation outside the blood vessels in between epithelial acini in pancreatitis (**Fig. 3B-C**).

**Figure 3.**
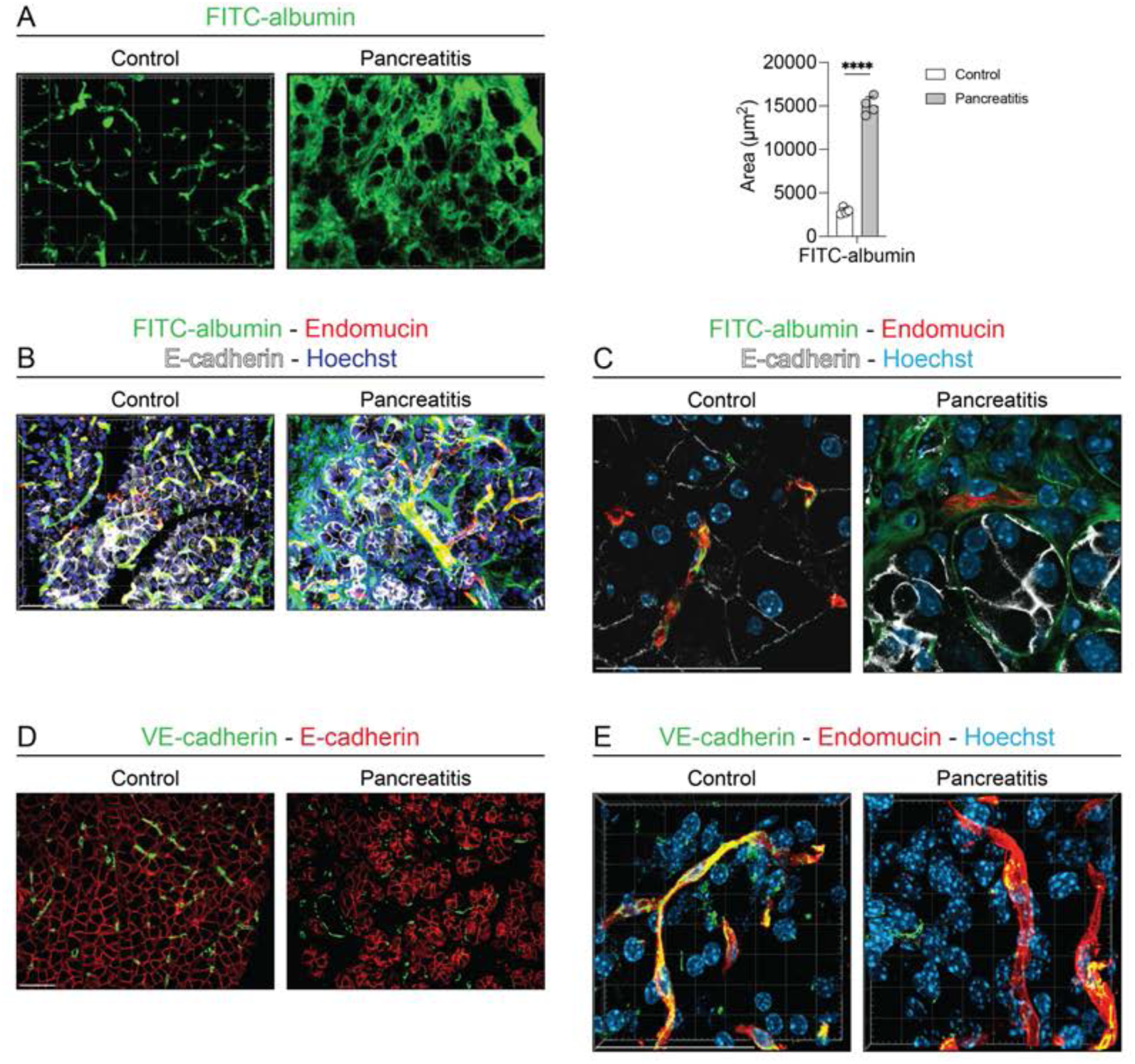
Blood vascular permeability during acute pancreatitis. (A) Visualization and quantification of FITC-albumin extravasation in pancreatic sections from control and caerulein-treated (acute pancreatitis) mice. Scale bar: 50 µm. Total area of FITC-albumin was quantified in a representative 40× focal plane. Data are shown as means ± SD. Statistical analysis: two-tailed Student’s t-test. \**p* < 0.05, \*\**p* < 0.01, \*\*\**p* < 0.001, \*\*\*\**p* < 0.0001. n=4. (B-C) Immunofluorescence labeling of endomucin (endothelial cells) and E-cadherin (epithelial cells), with nuclear counterstain (Hoechst) and FITC-albumin visualization in pancreatic sections from control and caerulein-treated. Low (B) and high (C) magnification are shown. Scale bar: 50 µm. (D-E) Immunofluorescence labeling of VE-cadherin (endothelial adherens junctions), and E-cadherin (epithelial cells) in pancreatic sections from control and caerulein-treated mice. Low (D) and high (E, also presenting the Hoechst nuclear counterstain) magnification are shown. Scale bar: 50 µm.

To propose a molecular mechanism for the vascular leakage, we studied the endothelial barrier function and in particular VE-cadherin, a critical component of the endothelial adherens junctions. As compared to control pancreas, VE-cadherin immunolabeling was weaker in pancreatitis tissues (**Fig. 3D**), despite increased mRNA expression level (**Fig. 2G**; *Cdh5*). Z-stack super-resolution imaging revealed almost perfect colocalization of VE-cadherin with endomucin in control pancreas (**Fig. 3E**, at left). However, in acute pancreatitis, VE-cadherin distribution was disrupted, appearing as punctate intracellular structures instead of being evenly distributed along the cell surface (**Fig. 3E**, at right). The absence of VE-cadherin at the cell surface suggests destabilization of endothelial adherens junctions, likely due to endocytosis and internalization, which may contribute to the increased vascular permeability observed in pancreatitis (**Fig. 3E**).

In summary, acute pancreatitis is also associated with increased vascular permeability and disrupted VE-cadherin localization, indicating compromised vascular integrity.

### VEGF-A expression increases during acute pancreatitis within metaplastic cells

To understand the molecular mechanisms underlying these morphological, functional, and molecular changes, we focused on VEGF-A, the primary regulator of vascular responses. Indeed, VEGF-A plays a crucial role in multiple endothelial processes, including cell survival, angiogenesis, cell morphology, adhesion, and permeability, acting via its main receptors VEGFR-1 (*Flt*) and VEGFR-2 (*Kdr*) both highly induced upon pancreatitis (**Fig. 2G**). We first analyzed the expression pattern of VEGF-A in the adult pancreas subjected to pancreatitis. RT-qPCR analysis revealed that *Vegfa* mRNA expression was significantly increased in pancreatitis, as compared to controls (**Fig. 4A**). This increase started to be detectable as of Day 4 and was highly significative at Day 6 (**Supplementary Fig. 3A**). To visualize and identify the cell type(s) producing VEGF-A, we performed *in situ* hybridization, using RNAscope™. In control pancreas, *Vegfa* mRNA was mainly expressed by cells in the islets of Langerhans but also in some acinar cells of the pancreatic parenchyma (**Fig. 4B**, at left and magnifications). Upon pancreatitis, and as observed by RT-qPCR, the signal intensity of the *Vegfa* probe increased and was found in almost all the E-cadherin-positive epithelial cells, including in ductal-like cells undergoing acinar-to-ductal metaplasia (**Fig. 4B**, at right). Thus, upon induction of pancreatitis, *Vegfa* expression is upregulated in all the acinar cells.

**Figure 4.**
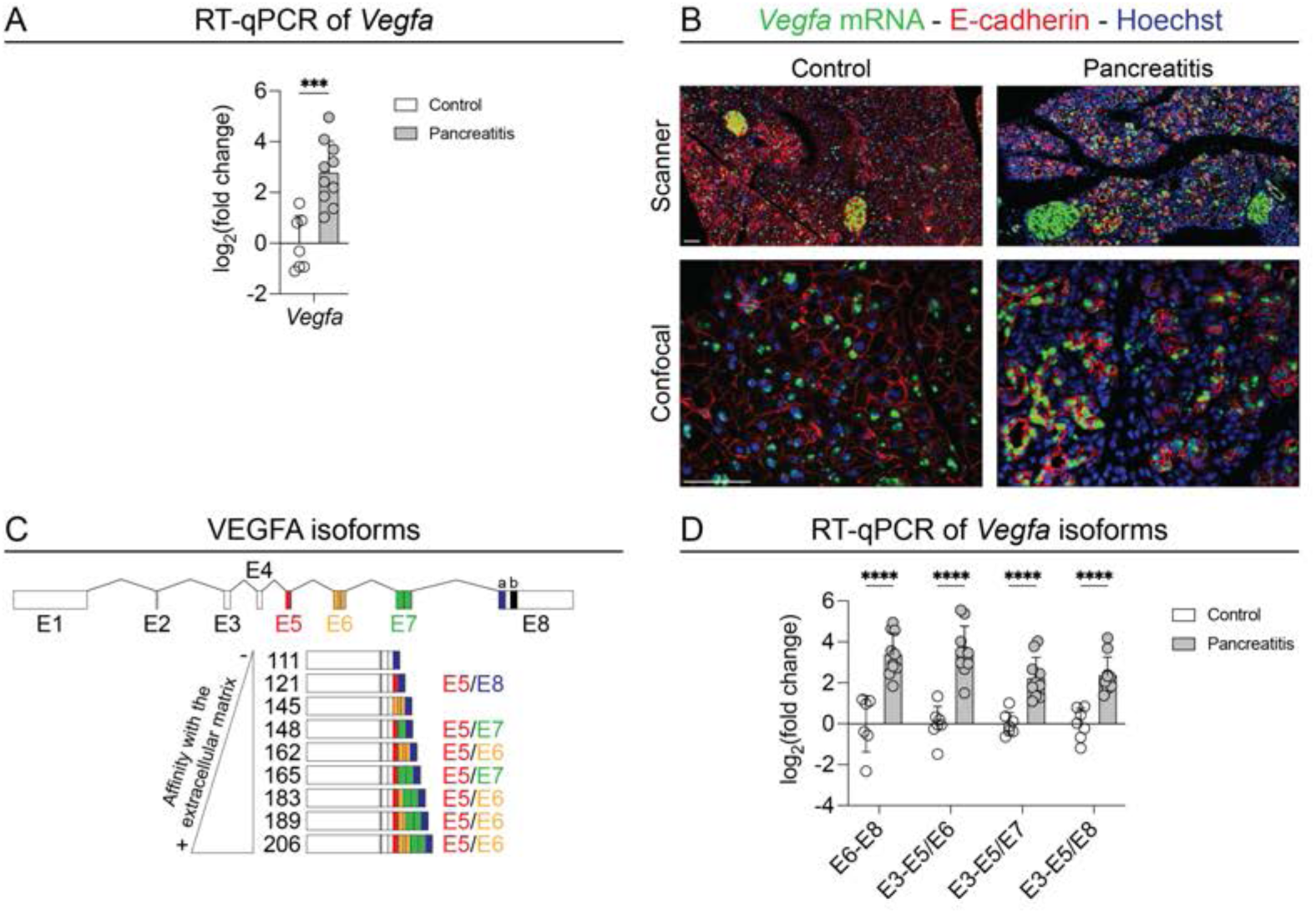
VEGF-A expression during acute pancreatitis. (A) Gene expression analysis by RT-qPCR of *Vegfa,* normalized to *18S* ribosomal RNA, in pancreatic tissue from control and caerulein-treated mice. Data are shown as means ± SD. Statistical analysis: two-tailed Student’s t-test. \**p* < 0.05, \*\**p* < 0.01, \*\*\**p* < 0.001, \*\*\*\**p* < 0.0001. n=7 for control, n=10 for pancreatitis. (B) *In situ* hybridization of *Vegfa* by RNAscope™ and immunofluorescence labeling of E-cadherin (epithelial cells), with a nuclear counterstain (Hoechst) in pancreatic sections from control and caerulein-treated mice. Scale bar: 50 µm. (C) Schematic representation of *Vegfa* gene organization with exons and introns and of VEGF-A splicing isoforms; the exon following exon 5 (E5) is shown at right. (D) Gene expression analysis by RT-qPCR of *Vegfa* splice variants (E6-E8, E3-E5/E6, E3-E5/E7, E3-E5/E8), normalized to *18S* ribosomal RNA, in pancreatic tissue from control and caerulein-treated mice. Data are shown as means ± SD. Statistical analysis: two-tailed Student’s t-test. \**p* < 0.05, \*\**p* < 0.01, \*\*\**p* < 0.001, \*\*\*\**p* < 0.0001. n≥6 for control, n=10 for pancreatitis.

VEGF-A exists in various isoforms due to alternative mRNA splicing. Exons 1 to 5 are common across most isoforms, while exons 6 and 7 encode heparin-binding regions, providing greater affinity for the extracellular matrix and consequently a more localized activity of this angiogenic factor. On the contrary, isoforms lacking exon 6 or exons 6 and 7 are more diffusible and are considered as long-acting VEGF-A isoforms (**Fig. 4C**). We therefore designed primer pairs to discriminate between VEGF-A isoforms based on exon retention. The E6-E8 and E3-E5/6 primer pairs detect locally-acting isoforms retaining respectively exons 6 and 7, and exon 6, while E3-E5/7 and E3-E5/8 primer pairs detect more diffusible isoforms lacking respectively exon 6, and exons 6 and 7. RT-qPCR analysis showed that all isoforms were upregulated in pancreatitis at Day 6, with a greater increase in locally acting isoforms retaining exon 6 (**Fig. 4D**, primers E6-E8 and E3-E5/6). Moreover, while the diffusible isoforms began to increase by Day 4, reaching statistical significance by Day 6, the locally acting isoforms showed an earlier increase, detectable by Day 2 and becoming significant by Day 4 (**Supplementary Fig. 3A**).

In conclusion, these findings demonstrate that VEGF-A expression increases during pancreatitis, particularly in acinar epithelial cells undergoing an acinar-to-ductal metaplasia. Locally acting isoforms showed a more important and rapid increase in expression, as compared to diffusible isoforms, suggesting that VEGF-A could have local effects near the metaplastic cells, which may be needed to adapt to the environmental stress.

### Acinar-specific *Vegfa* inactivation does not affect fibrosis, inflammation, and ADM during pancreatitis

To investigate the mechanisms by which acinar VEGF-A may drive vascular remodeling during acute pancreatitis, we specifically inactivated *Vegfa* in acinar cells. To this end, we crossed *Ptf1a-CreER^T2^* mice with *Vegfa^fl/fl^* mice to generate *Ptf1a-CreER^T2^; Vegfa^fl/fl^*. In these mice, the CreER^T2^ recombinase is specifically expressed in acinar cells via Ptf1a regulatory regions. Tamoxifen injections were used to trigger nuclear translocation of the CreER^T2^ and therefore the Cre-mediated recombination of *Vegfa* exon 3 (*Vegfa-E3*). This recombination causes a frameshift and leads to the specific knockout of VEGF-A in acinar cells (**Fig. 5A**). These knockout mice are called *Vegfa*^Δ*ac*^ and compared to *Vegfa^WT^* littermates (**Fig. 5A**).

**Figure 5.**
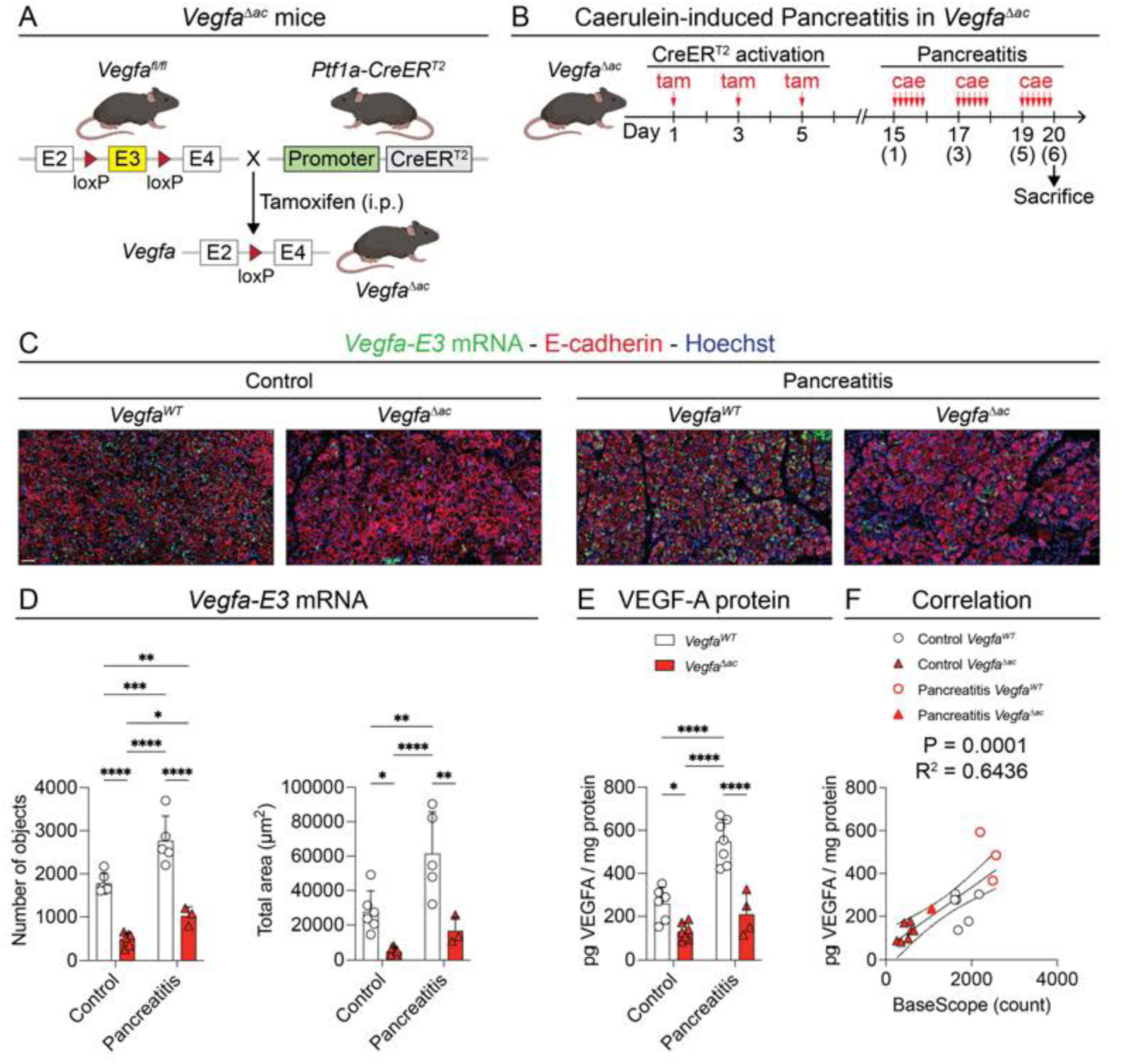
Acinar-specific inactivation of VEGF-A. (A) Schematic representation of the procedure for acinar-specific *Vegfa* inactivation using floxed *Vegfa* exon 3 allele (*Vegfa-E3*). CreER^T2^ expression is driven by the acinar-specific Ptf1a promoter and nuclear translocation by tamoxifen intraperitoneal injections (i.p.). (B) Timeline of tamoxifen and caerulein injections (vertical red arrows), and sacrifice for tissue harvest on day 6 (day 20 if counting from the initial tamoxifen injection) in *Vegfa^WT^* and *Vegfa*^Δ*ac*^ mice. (C) *In situ* hybridization of *Vegfa-E3* by BaseScope™ and immunofluorescence labeling of E-cadherin (epithelial cells), with a nuclear counterstain (Hoechst) in pancreatic sections from control and caerulein-treated *Vegfa^WT^* and *Vegfa*^Δ*ac*^ mice. Scale bar: 50 µm. (D) Quantification of *Vegfa-E3* objects: number and total area in pancreatic sections from control and caerulein-treated *Vegfa^WT^* and *Vegfa*^Δ*ac*^ mice. Data represent the average total area from one representative 488,160 µm^2^ field per mouse and are shown as means ± SD. Statistical analysis: two-way ANOVA with Holm-Šídák correction. \**p* < 0.05, \*\**p* < 0.01, \*\*\**p* < 0.001, \*\*\*\**p* < 0.0001. n≥3. (E) VEGF-A protein abundance measured by ELISA and normalized to total protein in pancreatic tissues from control and caerulein-treated *Vegfa^WT^* and *Vegfa*^Δ*ac*^ mice. Data are shown as means ± SD. Statistical analysis: two-way ANOVA with Holm-Šídák correction. \**p* < 0.05, \*\**p* < 0.01, \*\*\**p* < 0.001, \*\*\*\**p* < 0.0001. n≥4. (F) Correlation between ELISA measurements and BaseScope™ *Vegfa-E3* object counting, assessed using linear regression (straight middle line), with 95% confidence interval (curved lines around the middle line), Goodness of Fit (R^2^), and the significance of a non-zero slope (*p* value).

To induce the acinar-specific recombination, we administered tamoxifen three times over five days (**Fig. 5B**). Two weeks later, acute pancreatitis was induced using the same protocol as above (**Fig. 5B**; caerulein injection at Day 15 corresponds to injection at Day 1 in **Fig. 1A**). Mice body weight and pancreas weight remained unchanged in *Vegfa*^Δ*ac*^ mice, as compared to *Vegfa^WT^* littermates (**Supplementary Fig. 3E**).

To verify the acinar-specific recombination of *Vegfa*, we used a BaseScope™ *in situ* hybridization to visualize the exon 3 of *Vegfa* (*Vegfa-E3*). Although less sensitive, this exon 3 specific probe confirmed the increased expression in *Vegfa* mRNA during pancreatitis, particularly in epithelial metaplastic cells, consistent with results obtained using RNAscope™ (**Fig. 5C** to compare with **Fig. 4B**). Close-up images revealed the highest expression of *Vegfa* in islets of Langerhans and lower expression in acinar cells (**Supplementary Fig. 3B-C**). Interestingly, no *Vegfa* could be detected in ductal cells in healthy pancreas. Induction of pancreatitis induced the expression of *Vegfa* in the acinar cells but also in ductal cells (**Supplementary Fig. 3B-C**). As expected, in *Vegfa*^Δ*ac*^ mice, the *Vegfa-E3* probe hybridized to the islets of Langerhans and ductal structures but no signal was detected in the acinar compartment of the pancreatic parenchyma (**Fig. 5C** and **Supplementary Fig. 3B-C**). *Vegfa-E3* was also detected outside the pancreatic parenchyma during pancreatitis, likely reflecting *Vegfa* expression in infiltrating immune cells (**Fig. 5C**). Quantification of the number of objects and area positive for *Vegfa-E3* probe supported these observations and were highly correlated (**Fig. 5D** and **Supplementary Fig. 3D**). Additionally, ELISA measurements from total pancreas lysates confirmed increased VEGF-A protein levels during pancreatitis, and a reduction in *Vegfa*^Δ*ac*^ mice as compared to *Vegfa^WT^* (**Fig. 5E**). BaseScope and ELISA results displayed a good correlation (**Fig. 5F**).

We next evaluated the effects of VEGF-A acinar inactivation in untreated controls and during acute pancreatitis. Hematoxylin & eosin staining revealed no significant differences in tissue morphology between *Vegfa^WT^* and *Vegfa*^Δ*ac*^ mice, both in untreated controls and in acute pancreatitis (**Fig. 6A**). Similarly, immunofluorescence labeling of F4/80, vimentin, and E-cadherin revealed similar phenotype in control and pancreatitis samples of wild-type or knockout mice (**Fig. 6B-C**). Quantifications of the total labelled area and of ADM gene expression indicated that macrophage infiltration, fibrosis, or epithelial remodeling and ADM are independent from the acinar-specific increase in *Vegfa* observed during pancreatitis (**Fig. 6D-E**). Finally, the endocrine compartment, including the islets of Langerhans, also showed no significant changes in either genotype or condition (**Supplementary Fig. 3C**). To investigate the ability of pancreas to regenerate, pancreata were also harvested five days after the final caerulein injection (Day +5), instead of one day as in the previous experiments. In this context, both *Vegfa^WT^* and *Vegfa*^Δ*ac*^ mice showed similar regeneration, as visualized by the increase in eosinophilia and decreased macrophages (F4/80) and fibroblasts (Vimentin) labeling (**Supplementary Fig. 5A-D, F** to compare with **Fig. 6A-D**).

**Figure 6.**
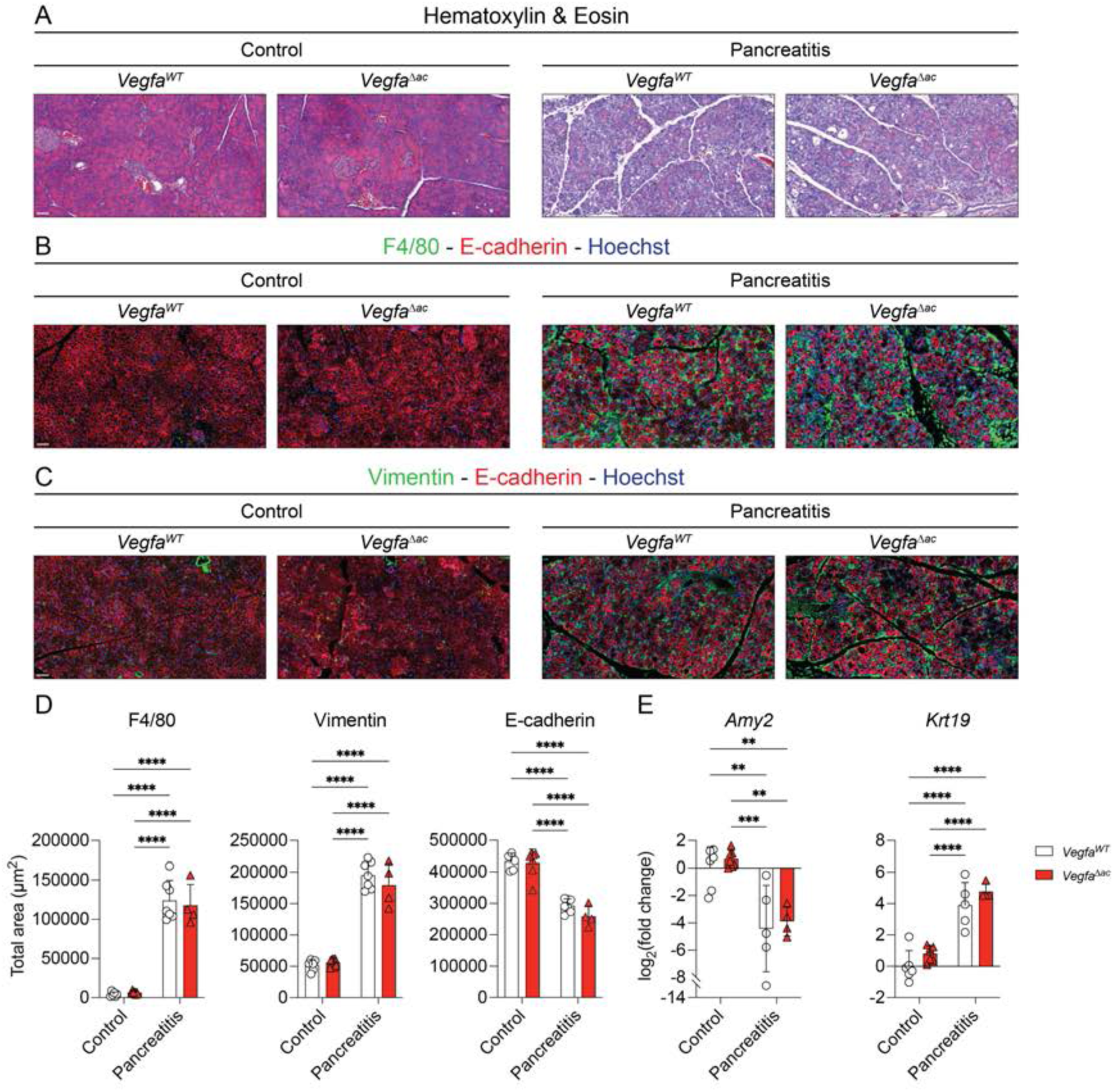
Effects of acinar-specific VEGF-A inactivation on acute pancreatitis. (A) Hematoxylin (nuclei) & eosin (cytoplasm) staining of pancreatic sections from control and caerulein-treated *Vegfa^WT^* and *Vegfa*^Δ*ac*^ mice. Scale bar: 50 µm. (B-C) Immunofluorescence labeling of F4/80 and E-cadherin (macrophages and epithelial cells; B) and of vimentin and E-cadherin (for fibroblasts and epithelial cells; C), with a nuclear counterstain (Hoechst) in pancreatic sections from control and caerulein-treated *Vegfa^WT^* and *Vegfa*^Δ*ac*^ mice. Scale bar: 50 µm. (D) Quantification of immunofluorescence labeling: total area (µm^2^) of F4/80, vimentin, and E-cadherin in pancreatic sections from control and caerulein-treated *Vegfa^WT^* and *Vegfa*^Δ*ac*^ mice. Data represent the average total area from three representative 488,160 µm^2^ fields per mouse and are shown as means ± SD. Statistical analysis: two-way ANOVA with Holm-Šídák correction. \**p* < 0.05, \*\**p* < 0.01, \*\*\**p* < 0.001, \*\*\*\**p* < 0.0001. n≥4. (E) Gene expression analysis by RT-qPCR of acinar (*Amy2*) and ductal (*Krt19*) markers, normalized to *18S* ribosomal RNA, in pancreatic tissue from control and caerulein-treated *Vegfa^WT^* and *Vegfa*^Δ*ac*^ mice. Data are shown as means ± SD. Statistical analysis: two-way ANOVA with Holm-Šídák correction. \**p* < 0.05, \*\**p* < 0.01, \*\*\**p* < 0.001, \*\*\*\**p* < 0.0001. n≥4.

In conclusion, these findings demonstrate that acinar-specific inactivation of VEGF-A does not affect fibrosis, inflammation, or acinar-to-ductal metaplasia during pancreatitis and its resolution, indicating that the increase in acinar VEGF-A observed during pancreatitis is not essential for these processes. This suggests that the vascular permeability driving immune cell infiltration and the inflammatory response in acute pancreatitis occurs independently of acinar VEGF-A.

### Acinar-specific *Vegfa* inactivation affects vascular remodeling during pancreatitis

We next examined whether the increased VEGF-A expression in acinar cells could control the vascular remodeling observed during acute pancreatitis. Immunofluorescence labeling of endomucin and of ERG, an endothelial-specific transcription factor was conducted (**Fig. 7A**), and endomucin-positive blood vessels were segmented across all four, genetic background and treatment, conditions (**Fig. 7B**).

**Figure 7.**
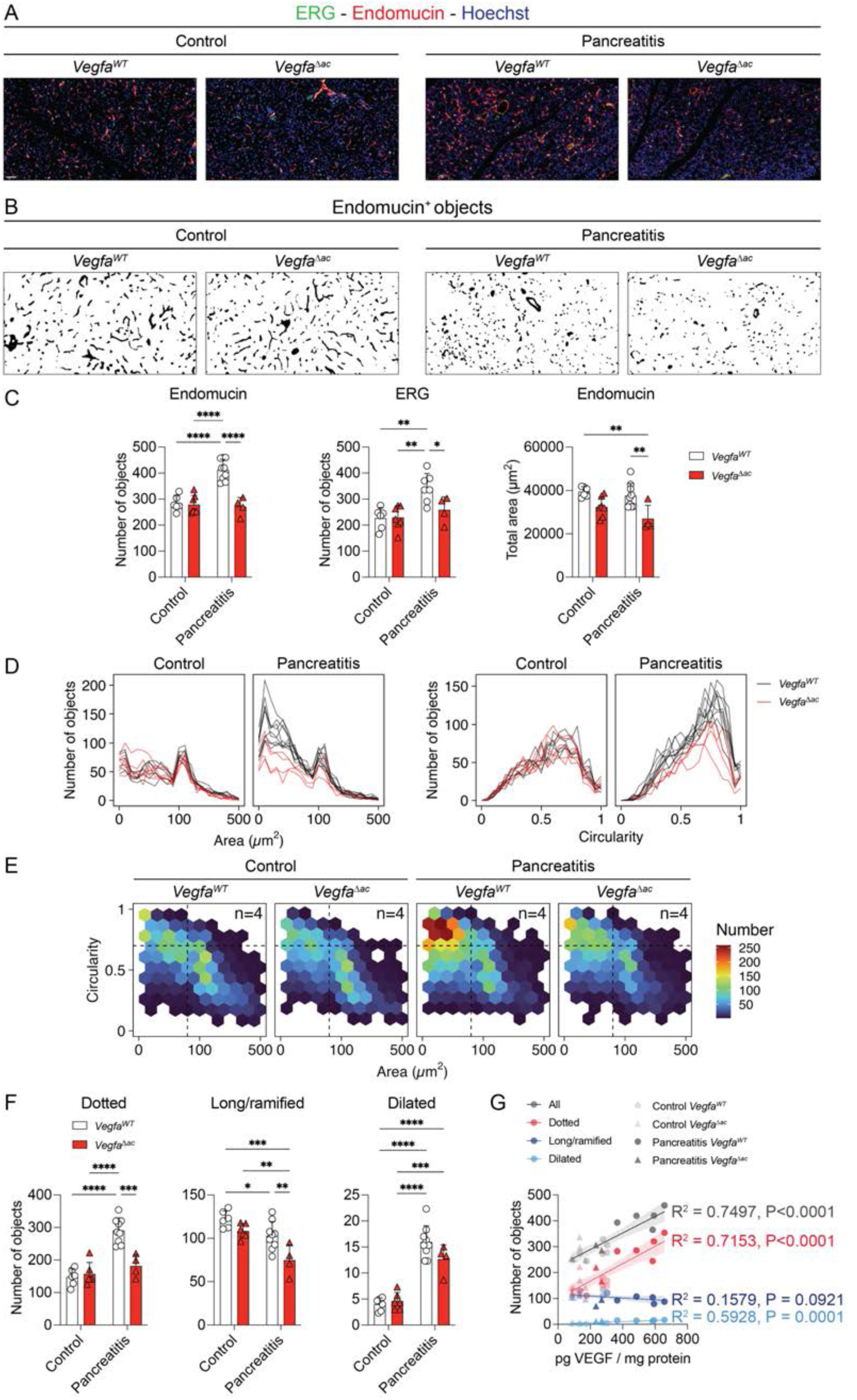
Effects of acinar-specific VEGF-A inactivation on blood vessels. (A) Immunofluorescence labeling of endomucin (endothelial cells) and ERG (endothelial cell nuclei), with a nuclear counterstain (Hoechst) in pancreatic sections from control and caerulein-treated *Vegfa^WT^* and *Vegfa*^Δ*ac*^ mice. Scale bar: 50 µm. (B) Segmentation of endomucin-positive objects from panel A for morphological analysis. (C) Quantification of blood vessel density: number of endomucin- and ERG-positive objects and total area of endomucin in pancreatic sections from control and caerulein-treated *Vegfa^WT^* and *Vegfa*^Δ*ac*^ mice. Data represent the average total area or counts from three representative 488,160 µm^2^ fields per mouse and are shown as means ± SD. Statistical analysis: two-way ANOVA with Holm-Šídák correction. \**p* < 0.05, \*\**p* < 0.01, \*\*\**p* < 0.001, \*\*\*\**p* < 0.0001. n≥4. (D) Quantification of endomucin-positive objects: area-based (at left) and circularity-based (at right) 1D count plot, in pancreatic sections from control and caerulein-treated *Vegfa^WT^* and *Vegfa*^Δ*ac*^ mice. Data represent the cumulative counts from three representative 488,160 µm^2^ fields per mouse. (E) Hexagonal binned plot of circularity vs. area (µm^2^) of endomucin-positive objects, with colors representing object count per bin in pancreatic sections from control and caerulein-treated *Vegfa^WT^* and *Vegfa*^Δ*ac*^ mice. Data represent the cumulative counts from three representative 488,160 µm^2^ fields per mouse. n=4. (F) Classification of endomucin-positive objects: dotted (area ≤ 80 µm^2^), long/ramified (area > 80 µm^2^, circularity < 0.7), and dilated (manually classified) in pancreatic sections from control and caerulein-treated *Vegfa^WT^* and *Vegfa*^Δ*ac*^ mice. Data represent the average counts from three representative 488,160 µm^2^ fields per mouse and are shown as means ± SD. Statistical analysis: two-tailed Student’s t-test. \**p* < 0.05, \*\**p* < 0.01, \*\*\**p* < 0.001, \*\*\*\**p* < 0.0001. n≥4. (G) Correlation between ELISA measurements and number of total objects (gray), dotted objects (red), long/ramified objects (blue), and dilated objects (light blue) was assessed using linear regression (straight middle line), with 95% confidence interval (colored area around the middle line), Goodness of Fit (R^2^), and the significance of a non-zero slope (*p* value).

Quantitative analysis confirmed the increase in blood vascular objects during pancreatitis. Specifically, the number of endomucin-positive and ERG-positive objects increased significantly in pancreatitis (**Fig. 7C**). Notably, this increase in endothelial objects in pancreatitis was suppressed in *Vegfa*^Δ*ac*^ mice compared to their *Vegfa^WT^* littermates, with values returning to levels observed in control pancreata (**Fig. 7C**). Furthermore, while the total area occupied by the endomucin signal was similar in control and caerulein-treated pancreata, it was significantly reduced in *Vegfa*^Δ*ac*^ mice (**Fig. 7C**). Acinar *Vegfa* inactivation specifically affected blood vessels within the acinar compartment, while vessels surrounding the ducts and those in the islets of Langerhans remained unaffected (**Supplementary Fig. 4**). Moreover, this effect on the vasculature persisted during regeneration at Day +5 (**Supplementary Fig. 5E–F**).

We also analyzed the specific characteristics of vascular remodeling during pancreatitis and how these were impacted by the loss of acinar VEGF-A. The high number of small vascular objects (< 80 µm^2^) and of circular objects (> 0.7) induced by pancreatitis in *Vegfa^WT^* mice was reduced in *Vegfa*^Δ*ac*^ mice (**Fig. 7D-E**), supporting the role of acinar VEGF-A in blood vessel remodeling. Finally, we analyzed if the number of dotted vascular objects and dilated vascular structures, which increased during pancreatitis, while long/ramified objects decreased (**Fig. 7F**, white bars), were affected in *Vegfa*^Δ*ac*^ pancreatitis mice (**Fig. 7F**, red bars). Interestingly, in *Vegfa*^Δ*ac*^, the number of dotted objects did not increase upon pancreatitis and remained at a level similar to untreated *Vegfa^WT^* mic e (**Fig. 7F**, at left). In addition, the decreased number of long/ramified objects was even more pronounced in *Vegfa*^Δ*ac*^ pancreatitis mice (**Fig. 7F**, middle). Lastly, the number of dilated structures increased to a slightly lower level in *Vegfa*^Δ*ac*^ mice, as compared to *Vegfa^WT^* mice during pancreatitis (**Fig. 7F**, at right). When plotting the number of vascular objects (all, dotted, long/ramified, or dilated) as a function of VEGF levels measured by ELISA, only the total number of vascular objects and the number of dotted objects correlated strongly with VEGF levels (**Fig. 7G**).

These results suggest that the increased expression of VEGF-A in acinar cells is critical for maintaining and preserving the vasculature. In the absence of VEGF-A secretion from acinar cells, the increase in vascular objects is impaired, resulting in fewer endothelial objects and reduced vessel surface area. These findings indicate that the vascular remodeling, and more specifically the increased number of dotted vascular objects induced upon pancreatitis depends on the acinar expression of VEGF-A. Overall, this demonstrates that VEGF-A is a key regulator of blood vessel adaptation and preservation during caerulein-induced pancreatitis.

## Discussion

In the present study, we focused on an essential but poorly studied compartment in pancreatitis, namely the blood vessels. We found significant blood vascular remodeling, with an increase in dotted and dilated vascular objects and a decrease in long-and-ramified vascular objects, as well as increased permeability after induction of acute pancreatitis in mice. Molecularly, pancreatitis was accompanied by increased expression of endothelial genes, and disrupted VE-cadherin distribution. Moreover, VEGF-A expression increased during pancreatitis. This increase was particularly evident in metaplastic acinar cells and concerned preferentially locally acting VEGF-A isoforms.

Using acinar-specific VEGF-A inactivation, we uncovered a key actor of the mechanism controlling vascular remodeling during pancreatitis. We also show that immune cell infiltration, fibrosis, and acinar-to-ductal metaplasia occur independently of VEGF-A, suggesting that vascular permeability, critical for the inflammatory response, may be independent of acinar VEGF-A.

### Pancreatitis is accompanied by changes of the microvasculature

Our findings reveal significant morphological remodeling of the blood vasculature during acute pancreatitis. Analysis of 2D sections revealed a marked reduction in long and ramified vascular objects, accompanied by an increase in dotted objects and dilated structures. In 3D, this remodeling was characterized by an apparent increase in vessel density and thickness, probably due to acinar tissue shrinking and vascular dilation. At the molecular level, these changes correlate with an upregulation of several endothelial markers, suggesting an active endothelial response during acute pancreatitis. We propose that this paradoxical pattern; i.e. thicker and denser vasculature in 3D, yet more dotted and less ramified in 2D; may represent pathological angiogenesis. Pathological angiogenesis is defined as the abnormal and often dysregulated growth of blood vessels in response to tissue injury. This process results in the formation of dysfunctional and disorganized vascular networks [25–27]. Key features of pathological angiogenesis include vessel dilation, endothelial activation, and the breakdown or restructuring of pre-existing vascular networks into immature and fragmented forms. These features align with the morphological alterations observed in our study. Within the inflammatory microenvironment of acute pancreatitis, endothelial cells may be activated in response to inflammatory signals, driving both vascular permeability and angiogenesis, ultimately contributing to the observed remodeling. The observed shift from long and ramified vessels to dotted structures could be explained by capillary tube regression, a phenomenon where elongated vessels lose their tubular integrity and collapse into disconnected or fragmented structures. Capillary regression can occur due to the action of matrix metalloproteinases and pro-inflammatory mediators, which degrade the extracellular matrix and destabilize vascular structures [28, 29]. In experimental models of pancreatitis, microvascular studies have similarly reported vessel fragmentation and irregularities in capillary walls, supporting this hypothesis [15, 16, 30, 31].

The increase in dilated vascular structures may reflect vasodilation, which could act as a compensatory response to the morphological alterations observed during acute inflammation. Acute pancreatitis is characterized by an early phase of vasoconstriction and capillary stasis, potentially explaining the increase in dotted objects, followed by a later phase of vasodilation and reperfusion, which may account for the observed rise in dilated vascular structures [32]. This dynamic process may lead to temporary vascular collapse and fragmentation during the early stages of the disease. This could also create a microenvironment conducive to tumor progression, potentially linking pancreatitis-associated vascular dysfunction to the hypovascularization of PDAC [24], accounting for the mechanism by which pancreatitis accelerates PDAC development [33].

Additionally, pancreatic intralobular arteriolar sphincter constriction and functional impairment have been shown to exacerbate microcirculatory disturbances [34], potentially contributing to the observed vascular changes. The increase in endothelial markers observed during acute pancreatitis in our study could also reflect endothelial activation, a hallmark of inflammation. Endothelial activation is commonly associated with vascular remodeling and permeability changes, and has been reported in pancreatic preinvasive lesions [11]. Although we did not study chronic pancreatitis, one could postulate that vascular remodeling upon repeated pancreatic injury could be maintained or exacerbated.

### Pancreatitis is accompanied by vascular permeability

The vascular leakage of FITC-albumin suggests increased vascular permeability, a phenomenon that has been identified in previous studies as an early and critical event in the pathophysiology of pancreatitis [18, 34–36]. The redistribution of VE-cadherin observed during acute pancreatitis has also been observed in experimental colitis, highlighting its involvement in inflammatory diseases [19]. Indeed, VE-cadherin plays a central role in maintaining vascular integrity, controlling vascular permeability, and regulating leukocyte extravasation [37–40]. Its disruption has been implicated in inflammation across various tissues, including the lung and skin [41]. Based on our findings, we propose that VE-cadherin internalization contributes to the vascular leakage observed in pancreatitis. By linking the redistribution of VE-cadherin to FITC-albumin leakage, we suggest that VE-cadherin is a key regulator of vascular permeability during acute pancreatitis.

### Metaplastic cells increase VEGF-A expression during acute pancreatitis

The increase in VEGF-A expression during acute pancreatitis has also been observed during inflammation of the colon and the lung [20, 21]. This suggests that VEGF-A may play a common role in the inflammatory response across various tissues, potentially driving vascular remodeling during inflammation in multiple organs.

Notably, all VEGF-A splice variants showed increased expression during acute pancreatitis, with the highest levels observed in variants that retain the matrix-binding exons 6 and 7. This may indicate that VEGF-A fulfills a dual role during acute pancreatitis: a predominant local function mediated by matrix-bound VEGF-A isoforms and a secondary, long-distance effect via diffusible isoforms. Indeed, the longer matrix-binding isoforms, such as VEGF-189 and VEGF-206, are known to be retained within the subepithelial extracellular matrix and exert chemotactic and chemokinetic effects, particularly on polymorphonuclear leukocytes [42, 43]. Conversely, the shorter, diffusible isoform VEGF-121 has been shown to preserve microvessels in the context of acute kidney injury [44]. Although the cellular source of these isoforms is unknown, the multiple roles of VEGF-A in acute pancreatitis may be splice variant-dependent, with longer isoforms contributing to localized chemotactic activity, while shorter ones preserving microvasculature.

The upregulation of VEGF-A in acinar cells during acute pancreatitis may be linked to ADM, a process in which acinar cells undergo significant phenotypic and transcriptional changes in response to inflammation. ADM is triggered by proinflammatory cytokines released by macrophages during pancreatitis [7]. Similar cytokine-driven VEGF-A upregulation has been reported in other contexts, including alveolar epithelial cells under inflammatory conditions [45], and cervical cancer, where IL-6 activates STAT3 to drive VEGF-A expression [46]. The transcriptional reprogramming associated with ADM may also contribute to increased VEGF-A expression. Pathways such as NF-kB, Notch, and JAK-STAT3 are highly activated during ADM and may be implicated in VEGF-A regulation [47–49, 7, 50]. STAT3 activation, in particular, has been shown to upregulate VEGF-A and promote angiogenesis in various tumors [46, 51]. These findings suggest that inflammatory cytokines, especially IL-6, may upregulate VEGF-A expression in metaplastic acinar cells through a STAT3-dependent mechanism during acute pancreatitis. Alternatively, increased VEGF-A expression might be due to the ductal-like phenotype. Indeed, during development, VEGF-A is highly expressed by ductal progenitors and much less by acinar cells [52]. This regionalized expression of VEGF-A is key to recruit endothelial cells around ductal and endocrine progenitors and much less around developing acinar structures.

Given that acinar cells express VEGF-A in the control pancreas, it is possible that acinar cell death preceding inflammation leads to the release of VEGF-A like a damage-associated molecular pattern (DAMP). This early VEGF-A signaling could be important in triggering initial inflammation, though it may not play a role once the inflammatory response is fully amplified. Investigating VEGF-A dynamics at very early timepoints could help determine its contribution to the onset of pancreatitis.

### Acinar VEGF-A affects blood vessels during acute pancreatitis

In our study, we specifically targeted VEGF-A from acinar cells, the predominant epithelial cell type in the pancreas, where VEGF-A was most notably upregulated during pancreatitis. Acinar VEGF-A inactivation had a vascular effect but did not influence immune cell infiltration, ADM, or fibrosis. These findings suggest that acinar-derived VEGF-A primarily regulates vascular remodeling during pancreatitis, without affecting leukocyte infiltration and vascular permeability. While our ELISA and BaseScope™ analyses confirmed that acinar-specific VEGF-A inactivation significantly reduced VEGF-A levels during pancreatitis, residual VEGF-A was still detectable in knockout mice, suggesting contributions from non-acinar sources.

It is possible that acinar and non-acinar VEGF-A have distinct roles in acute pancreatitis. Acinar-derived VEGF-A appears critical for microvessel preservation, possibly through preferential expression of short-diffusible VEGF-121, as observed in acute kidney injury [44]. In contrast, non-acinar VEGF-A may contribute to pro-inflammatory processes, including leukocyte extravasation, vascular leakage, and permeability. Previous studies have demonstrated that VEGF-A mediates vascular permeability via VE-cadherin endocytosis [53], and regulates both angiogenesis and inflammation in inflammatory bowel disease [54]. Neutralizing VEGF-A in models of ulcerative colitis and sepsis-associated acute lung injury was also shown to reduce vascular permeability, leukocyte infiltration, and disease severity [19–21]. Among the non-acinar sources of VEGF-A, macrophages, a prominent cell type in acute pancreatitis [7], have been shown to produce VEGF-A [55, 56]. Furthermore, macrophage-derived VEGF-A can regulate inflammation through non-angiogenic mechanisms, such as during mycobacterial infection [57]. So, while our study highlights the role of acinar VEGF-A in preserving vascular integrity during acute pancreatitis, it is plausible that non-acinar VEGF-A, particularly from macrophages, plays a complementary role in promoting inflammation and vascular permeability. Further studies are needed to define the distinct contributions of VEGF-A from non-acinar sources, as well as the specific roles of its isoforms, in the pathophysiology of pancreatitis. To explore the role of non-acinar VEGF-A, we could selectively inactivate *Vegfa* in myeloid cells (*Vegfa*^Δ*mye*^) using *LysM-Cre; Vegfa^fl/fl^* mice, in fibroblasts (*Vegfa*^Δ*fib*^) using *Col1a2-Cre; Vegfa^fl/fl^* mice, and in ductal cells (*Vegfa*^Δ*duct*^) using *Sox9-CreER; Vegfa^fl/fl^* mice. In parallel, to investigate isoform-specific functions, particularly of matrix-bound VEGF-A, we could use floxed *Vegfa-E6* and/or *Vegfa-E7* alleles, instead of *Vegfa-E3*.

Besides VEGF-A, other mediators could contribute to vascular permeability during pancreatitis, potentially compensating for the effects of acinar VEGF-A. For instance, endothelins and bradykinin have been implicated in vascular regulation during inflammation. Endothelin 1 (ET-1) has been shown to alter pancreatic microcirculation and reduce blood flow during pancreatitis [58, 59]. Similarly, bradykinin, a peptide known to promote inflammation by inducing vasodilation and increasing vascular permeability, has been demonstrated to affect pancreatitis. Inhibition of the bradykinin B2 receptor preserved microcirculation in experimental pancreatitis models in rats [17]. These findings suggest that multiple mediators work in concert to regulate blood vessels during pancreatitis, potentially overshadowing the role of acinar-derived VEGF-A. While VEGF-A was selected based on its well-established role in vascular biology, an unbiased approach, such as transcriptomic or ligand-receptor interaction analyses, could provide a more comprehensive understanding of the mediators involved. Such methodologies have been applied to study pancreas development [60], and could be valuable in identifying additional regulators of blood vessels during pancreatitis.

In our study, acinar VEGF-A specifically played a role in acute pancreatitis but was dispensable in the normal pancreas. This aligns with a previous study demonstrating that epithelial VEGF-A, derived from acinar, ductal, and endocrine cells (islets of Langerhans), is not required for maintaining the vasculature of the adult pancreas under homeostatic conditions [61]. Mature blood vessels in the adult pancreas likely do not depend on VEGF-A for survival. In contrast, blood vessels associated with pancreatitis, particularly those appearing in 2D as dotted vessels in our study, appear immature and more dependent on acinar-derived VEGF-A. This phenomenon mirrors observations in xenografted gliomas, where immature vessels were selectively vulnerable to VEGF-A loss, unlike mature vessels [22]. Authors attributed this vulnerability to the absence of periendothelial cells, which normally provide stability and support to mature vessels [22].

The role of acinar VEGF-A during acute pancreatitis also parallels its function during pancreatic development. Inactivation of epithelial VEGF-A during development has been shown to reduce endothelial cell density [52]. In our study, VEGF-A was specifically inactivated in acinar cells, which undergo ADM during pancreatitis. ADM is often viewed not only as a transdifferentiation process but also as a dedifferentiation of acinar cells into a progenitor-like state. Unlike normal ductal cells, metaplastic ductal-like cells are proliferative, can revert to the acinar cell state, and contribute to pancreatic regeneration after injury [62]. By reactivating progenitor-associated genes, acinar cells undergoing ADM may also reestablish developmental acinar-to-endothelial interactions, which could be critical for tissue healing and regeneration following pancreatic injury [60, 63].

## Methods

### Ethics statement

All mice were housed in individually ventilated cages (IVC) and handled in accordance with the NIH Guide for the Care and Use of Laboratory Animals. Experiments were approved by the UCLouvain Animal Ethics Committee (2020/UCL/MD/011 and 2023/UCL/MD/47) and complied with ARRIVE guidelines.

### Mice and treatments

Wild-type C57BL/6N mice were purchased from Charles River Laboratories. Tg(*Ptf1a^Cre-ERTM^*) were kindly gifted by Francisco X. Real [64]. Tg(*Vegfa^tm2Gne^*) mice were obtained from Napoleone Ferrara (Genentech) [65]. Transgenic mice were generated by crossing male hemizygous Tg(*Ptf1a^Cre-ERTM^*) mice, hereafter referred to as *Ptf1a-CreER^T2^* [64], with female Tg(*Vegfa^tm2Gne^*) mice, hereafter referred to as *Vegfa^fl/fl^*[65]. Matings produced *Vegfa*^Δ*ac*^ mice (*Ptf1a-CreER^T2^; Vegfa^fl/fl^*), which were compared to their wild-type littermates (*Vegfa^WT^*), matching in age, and not carrying the *Ptf1a-CreER^T2^* transgene. All mice were on a C57BL/6N genetic background.

In *Vegfa*^Δ*ac*^ mice, acinar-specific recombination of *Vegfa* was induced using tamoxifen injections. Tamoxifen (T5648, Sigma-Aldrich) was dissolved in corn oil at a concentration of 25 mg/mL and administered intraperitoneally to both *Vegfa*^Δ*ac*^ and *Vegfa^WT^* mice at 6-8 weeks of age. The dosage was 125 µg/g body weight, given every 48 hours over five days (on days 1, 3, and 5). After the final injection on day 5, mice underwent a one-week rest before further experiments.

Acute pancreatitis was induced in wild-type C57BL/6N mice aged 6-8 weeks or in transgenic *Vegfa*^Δ*ac*^ and *Vegfa^WT^* mice after the tamoxifen rest week. Caerulein (AS-24252, Eurogentec) was dissolved in sterile phosphate-buffered saline (PBS) at a concentration of 37.5µg/mL. Mice received six hourly intraperitoneal injections of caerulein at a dosage of 125µg/kg body weight, administered every 48 hours over five days. The pancreas was harvested on day 2, day 4, the day after the final caerulein injection (day 6), or five days after the final injection (day 10).

For vascular permeability analysis, FITC-albumin was administered on day 6, 10 minutes before sacrifice. Mice were anesthetized with ketamine/xylazine and FITC-albumin or albumin–fluorescein isothiocyanate conjugate (A9771, Sigma-Aldrich) was injected into the vena cava. The FITC-albumin solution was prepared at a concentration of 70 mg/mL in sterile physiological saline (0.9% sodium chloride) and administered at a dosage of 210 µg/g body weight.

### Histology, mRNA *in situ* hybridization, immunofluorescence, and microscopy

#### Tissue sections

Dissected pancreata were fixed in 4% paraformaldehyde in PBS at 4°C overnight. Tissue samples were either prepared for paraffin sections or for cryosections, depending on experimental requirements. For paraffin sections (6µm), the pancreas was dehydrated and embedded in paraffin using a Tissue-Tek VIP-6 (Sakura), then sectioned at 6µm. For cryosections, necessary to preserve FITC-albumin fluorescence, tissues were equilibrated in 20% sucrose at 4°C overnight, embedded in 7.5% gelatin/15% sucrose, frozen at −80°C, and sectioned at 20µm.

#### Histological staining

Deparaffinized sections were stained with hematoxylin and eosin (H&E) or sirius red/fast green. Hematoxylin stains nucleic acids, highlighting the nuclei, while eosin stains proteins found in the cytoplasm and extracellular matrix. Sirius red binds to collagen, while fast green stains the remaining proteins, particularly those in the cytoplasm.

#### mRNA *in situ* hybridization

Paraffin sections were deparaffinized and processed for mRNA *in situ* hybridization using either the RNAscope™ 2.5 HD Assay - RED or the BaseScope™ RED Assay, following the manufacturer’s instructions. RNAscope™ was conducted to detect whole *Vegfa* mRNA with the Mm-Vegfa-36zz-C1 probe (31293-C1). BaseScope™ was used for the specific detection of *Vegfa* exon 3, using the BA-Mm-Vegfa-E3-3zz-st-C1 probe (1238051-C1). Following the mRNA *in situ* hybridization protocol, slides were subjected to immunofluorescence colabeling, beginning with the blocking step.

#### Immunofluorescence

Paraffin sections were deparaffinized and subjected to antigen retrieval by boiling twice for 5 minutes in 10mM sodium citrate (pH 6.0) in a 750W microwave. FITC-albumin cryosections were fixed in 4% paraformaldehyde in PBS for 15 minutes at room temperature, followed by heating in 40°C PBS for 5 minutes to remove gelatin. Sections were permeabilized with 0.3% Triton X-100 in PBS for 5 minutes at room temperature. Blocking was performed in 3% milk, 10% bovine serum albumin (BSA), and 0.3% Triton X-100 in PBS for 45 minutes at room temperature. Slides were incubated at 4°C overnight with primary antibodies diluted in blocking buffer. The following day, sections were washed and incubated at room temperature for 1 hour with 250 ng/mL Hoechst 33342 nuclear counterstain and secondary antibodies diluted 1:500 in blocking buffer without milk.

#### 3D imaging

Dissected pancreas tissues were fixed in 4% PFA in PBS at 4°C overnight. After fixation, the tissues were transferred to TBST (50 mM Tris HCl pH 7.5, 150 mM NaCl, 0.1% Triton X-100) and blocked in TBST containing 10% normal goat serum for 45 minutes at room temperature. The samples were incubated at 4°C overnight with primary antibodies diluted in blocking buffer, followed by at least five thorough 1-hour washes in TBST. Tissues were then incubated at 4°C overnight with secondary antibodies diluted in blocking buffer. After additional TBST washes (at least five 1-hour washes), the pancreas tissues were post-fixed in 4% PFA for 10 minutes at room temperature and stored in TBST until imaging.

#### Antibodies

The following primary antibodies were used: anti-E-cadherin (dilution 1:250, mouse IgG2a, 610182, BD), anti-vimentin (dilution 1:100, rabbit, 5741, Cell Signaling), anti-F4/80 (dilution 1:250, rabbit, 70076, Cell Signaling), anti-endomucin (dilution 1:100, rat, sc-65495, Santa Cruz), anti-ERG (dilution 1:500, rabbit, ab92513, Abcam), anti-VE-cadherin (dilution 1:100, goat, AF1002, R&D). The following secondary antibodies were used: Goat anti-Rat IgG (Alexa Fluor™ 647, A21247, Invitrogen), Goat anti-Mouse IgG2a (Alexa Fluor™ 568, A21134, Invitrogen), Goat anti-Rabbit IgG (Alexa Fluor™ 488, A11034, Invitrogen), Donkey anti-Goat IgG (Alexa Fluor™ 647, A21447, Invitrogen), Donkey anti-Rat IgG (Alexa Fluor™ 594, A21209, Invitrogen), Donkey anti-Mouse IgG (Alexa Fluor™ 488, A21202, Invitrogen).

#### Mounting

Slides were mounted using Dako Faramount Aqueous Mounting Medium (Agilent Technologies, Santa Clara, CA, USA).

#### Microscopy and imaging

Stained or immunolabelled sections were imaged using a Pannoramic 250 Flash III digital slide scanner (3DHistech). Cryosections (FITC-albumin) were visualized using a Zeiss Cell Observer Spinning Disk confocal microscope, using a 40× objective. For super-resolution visualization of FITC-albumin distribution and VE-cadherin localization, a Zeiss LSM 980 equipped with Airyscan 2 was employed. Imaging was performed using a Plan-Apochromat 63×/1.4 Oil DIC M27 Objective. Confocal microscopy images were either acquired at a single X-Y focal plane or as a Z-stack (X-Y-Z).

### Image analysis and quantification

Quantification was primarily conducted on slides scanned with the Pannoramic 250 Flash III digital slide scanner (3DHistech), except for FITC-albumin, which was analyzed on single representative 40× focal planes imaged using the Zeiss Cell Observer Spinning Disk confocal microscope.

For the quantification of F4/80, vimentin, and endomucin, at least three randomly selected fields, at 20× magnification, were analyzed per mouse pancreatic tissue section. These fields, selected with the CaseViewer software (3DHistech), corresponded to a surface of 488,160 µm^2^ each, resulting in a total quantified surface of 1,464,480 µm^2^ per mouse. *Vegfa-E3* expression was quantified using BaseScope™ on a representative 488,160 µm^2^ field per mouse.

Quantification was performed using ImageJ software. When necessary, image channels were split, and the resulting images were converted to 8-bit grayscale. Thresholding, a segmentation process that converts grayscale images into black-and-white binary images, was applied to distinguish the signal (white) from the background (black). The binary images were then analyzed to quantify signal particles. For each field, a summary report was generated that included the total area (µm^2^) occupied by the signal and the count of objects per section. For endomucin-positive objects, additional shape descriptors were analyzed at both the individual object level and across fields. These included 1) the area (µm^2^), representing the surface occupied by each object; 2) circularity, calculated as 4π × area/perimeter^2^, where a value of 1 indicates a perfect circle and values approaching 0 indicate elongated shapes; 3) the aspect ratio, defined as the ratio of the major axis to the minor axis of an ellipse fitted to the object, with higher values indicating greater elongation; and 4) roundness, calculated as 4 × area / (π × major axis^2^), which is essentially the inverse of the aspect ratio. Like circularity, roundness reflects how closely an object resembles a circle, with values closer to 1 indicating rounder shapes.

1D-density plots were created by plotting the number of objects (y-axis) against the analyzed parameter (x-axis). To visualize the relationship between circularity and area, 2D-density plots were generated, where each bin was color-coded to represent the number of objects within that range. Further categorization of endomucin-positive objects into dotted objects, long/ramified objects, and dilated structures was performed following these steps. First, dilated structures, characterized by the presence of openings, were manually identified, separated, and counted. Next, the remaining non-dilated objects were classified based on their area and circularity: dotted objects were defined as having an area ≤ 80 µm^2^, while long/ramified objects were defined as having an area > 80 µm^2^ and a circularity < 0.7. The thresholds were chosen based on empirical observations. The thresholds were chosen based on empirical observations. The area threshold of 80 µm^2^ was used to distinguish small, punctate structures from larger endothelial objects. The circularity threshold of 0.7 was selected to differentiate long/ramified objects from more circular structures.

### RNA isolation and RT-qPCR

Pancreatic tissue samples, measuring approximately 2 × 2 mm, were microdissected, immediately submerged in RNAlater™ Stabilization Solution (AM7021, Invitrogen) to preserve RNA integrity, and snap-frozen in liquid nitrogen. Total RNA was extracted using the ReliaPrep™ RNA Miniprep System (Z6111, Promega), following the manufacturer’s protocol optimized for small fibrous tissue samples (≤ 5 mg), as previously described [66]. Reverse transcription was performed using 500 ng of total RNA. The reaction was carried out with M-MLV Reverse Transcriptase (28025013, Invitrogen) and random hexamers. Quantitative PCR was conducted using the resulting cDNA, the KAPA SYBR® FAST qPCR Master Mix (2X) (KK4602, Roche), and target cDNA-specific primer pairs. Gene expression levels were analyzed using the 2^-ΔΔCt^ method, with *18S* ribosomal RNA serving as the reference RNA. Fold changes in gene expression were calculated, and the log2-transformed values of these fold changes were presented in the graphs.

The following primer pairs were used: 5’-GTAACCCGTTGAACCCCATT-3’ and 5’- CCATCCAATCGGTAGTAGCG-3’ for *18S*, 5’-CTCCTGACAAGGAGGAGCTG-3’ and 5’-ATAGTGCTCCCACTGGCTTG -3’ for *Cpa1*, 5’-TTCTGATCCTAGCCCTTGTG-3’ and 5’-TGATAGGGGACAGAACTCTC-3’ for *Prss2*, 5’- GTGGTCAATGGTCAGCCTTT-3’ and 5’- TTGCCATCGACCTTATCTCC-3’ for *Amy2*, 5’-ACCCTCCCGAGATTACAACC-3’ and 5’- GGCGAGCATTGTCAATCTGT-3’ for *Krt19*, 5’-CAAGACTCTGGGCAAGCTCTG-3’ and 5’- TCCGCTTGTCCGTTCTTCAC- 3’ for *Sox9*, 5’-GGTACTAGTCCGTGGTTCTTC-3’ and 5’- TTCCAGCGCATGTCGGCGCTC -3’ for *Onecut1*, 5’-GGCTTTGCAGATCCACATTT-3’ and 5’-GAGCCCAGCAGAGTGCTAGT-3’ for *Flt1*, 5’-GCATGGAAGAGGATTCTGGA- 3’ and 5’-CGGCTCTTTCGCTTACTGTT-3’ for *Kdr*, 5’-GGATGTGGTGCCAGTAAACC-3’ and 5’-ACCCCGTTGTCTGAGATGAG-3’ for *Cdh5*, 5’- ATAGGCATCAGCTGCCAGTC-3’ and 5’-TCCGCTCTGCACTGGTATTC-3’ for *Pecam1*, 5’-AATCAGGGCGCTATGGCTTT-3’ and 5’- TACTGAGCAAGATGCTGGGCAA-3’ for *Sox18*, 5’-TGACCACAAATGAGCGCAGA-3’ and 5’-GCCATATTCTTTCACCGCCC-3’ for *Erg*, 5’-CTGCATGGTGATGTTGCTCT-3’ and 5’-GTACCTCCACCATGCCAAGT-3’ for *Vegfa*, 5’-AATCCCGGTTTAAATCCTGG- 3’ and 5’-CACCGCCTTGGCTTGTCAC-3’ for *Vegfa E6-E8*, 5’- TGAGACCCTGGTGGACATCT-3’ and 5’-CTTTCCTCGAACTGATTTTTTTTC-3’ for *Vegfa E3-E5/E6*, 5’-TGAGACCCTGGTGGACATCT-3’ and 5’- TGAACAAGGCTCACAGTGATTTTC-3’ for *Vegfa E3-E5/E7*, 5’- TGAGACCCTGGTGGACATCT-3’ and 5’-GCCTTGGCTTGTCACATTTTTC-3’ for *Vegfa E3-E5/E8*.

### ELISA

Pancreatic tissue sample was dissected and lysed in 10µL of classical RIPA buffer (without SDS) per mg of tissue. The buffer was supplemented with cOmplete™ Mini EDTA-free Protease Inhibitor Cocktail (11836170001, Roche). Tissue homogenization was performed using a TissueLyser LT (Qiagen). Total protein concentration was determined via a bicinchoninic acid (BCA) assay. The samples were then diluted 1:10, and VEGF-A concentration was quantified using the Mouse VEGF ELISA Kit - Quantikine (MMV00, bio-techne), according to the manufacturer’s instructions. VEGF- A levels (pg/mL) were normalized to total protein concentration (mg/mL) and presented as pg of VEGF-A per mg of total protein in the graphs.

### Statistical analysis

GraphPad Prism and RStudio were used for data presentation, analysis, and statistical testing. All density plots were generated using RStudio. Statistical analyses included a two-tailed Student’s t-test for comparisons between two groups and a two-way ANOVA for comparisons across four groups. The normality of sample distributions was assessed using the D’Agostino-Pearson omnibus (K2) test and the Shapiro-Wilk test, while Spearman’s test for heteroscedasticity was used to evaluate sample homoscedasticity. For multiple comparisons, the Holm-Šídák method was applied. A *p* value of < 0.05 was considered statistically significant.

## Author Contributions

Conceptualization: E.A. and C.E.P.; formal analysis: E.A. and C.E.P.; investigation: E.A., H.L., S.M., D.F., L.V.; resources: P.H., D.T. and C.E.P.; writing—original draft preparation: E.A. and C.E.P.; supervision: C.E.P.; funding acquisition: E.A. and C.E.P. All authors have read and agreed to the published version of the manuscript. Authors thank Genentech for the Vegfa-floxed mice and Francisco X. Real for the *Ptf1a-CreER^T2^* mice.

## Writing Assistance

E.A. holds a fellowship from the Fonds pour la Formation à la Recherche dans l’Industrie et dans l’Agriculture (FRIA, Belgium), H.L. holds a Télévie fellowship (FNRS, Belgium), D.F. is a research assistant from UCLouvain, D.T. is a Senior Research Associate at the F.R.S.-FNRS.

## Grant Support

This study was supported by grants from the Université catholique de Louvain and F.R.S.-FNRS (Application ID: 40020690).

## Abbreviations

VEGF-A: (Vascular Endothelial Growth Factor A)
ADM: (acinar-to-ductal metaplasia)
PDAC: (pancreatic ductal adenocarcinoma)
RT-qPCR: (quantitative reverse transcription polymerase chain reaction)
FITC-albumin: (fluorescein isothiocyanate-tagged albumin)
VEGFR: (vascular endothelial growth factor receptor)
ELISA: (enzyme-linked immunosorbent assay)
SD: (standard deviation).

## Data Transparency

The data, analytic methods, and study materials used in this manuscript will be made available to other researchers upon reasonable request.

## Statements and declarations

All authors confirm that they have no conflicts of interest relevant to this manuscript.

## Supplementary Figures

**Supplementary Figure 1.**
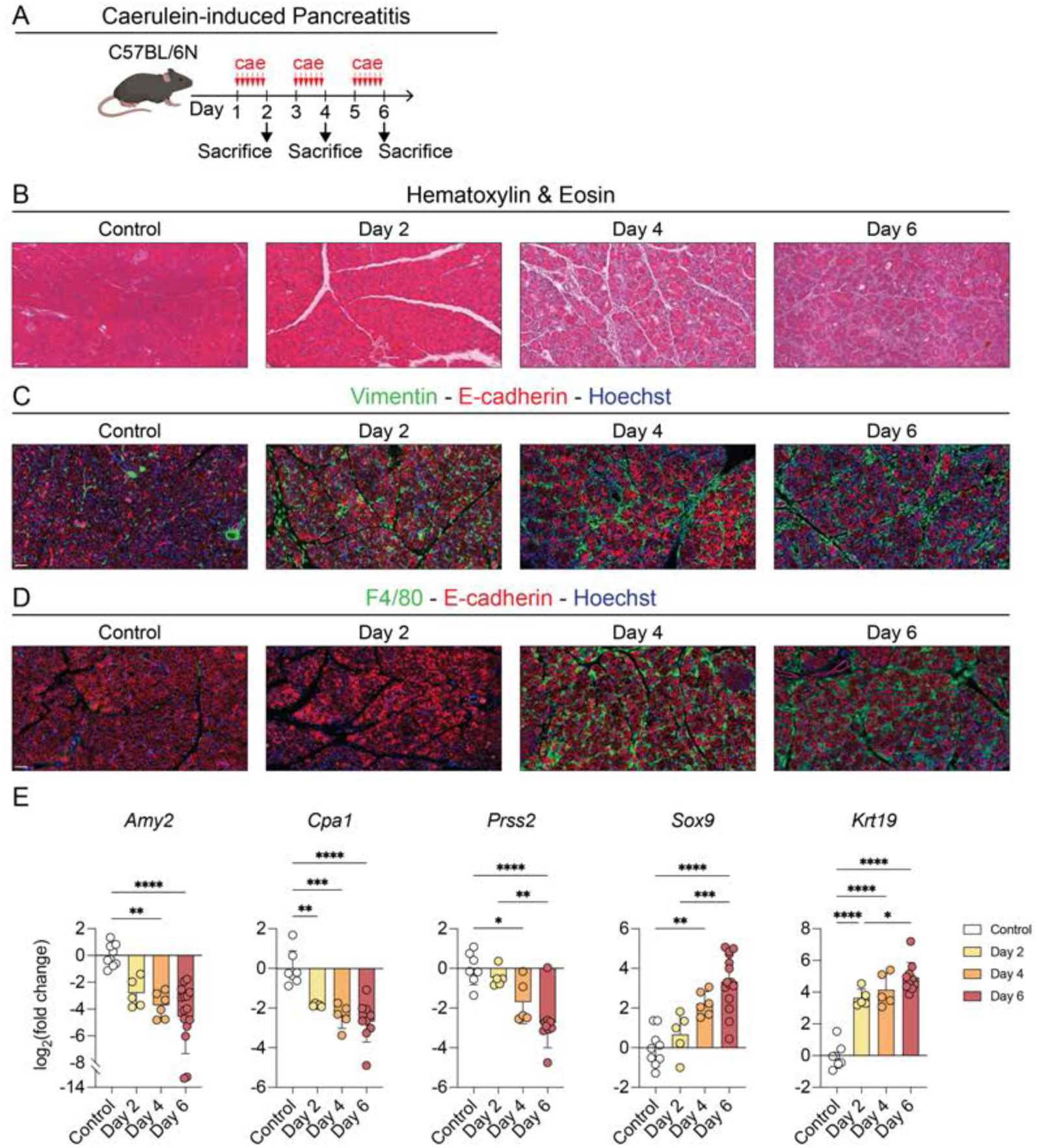
Time-course of acute pancreatitis. (A) Timeline of caerulein injections and sacrifice for tissue harvest on days 2, 4, and 6 in C57BL/6N mice. Vertical red arrows illustrate the six hourly injections. (B) Hematoxylin (nuclei) & eosin (cytoplasm) staining of pancreatic sections from control, day 2, day 4, and day 6 mice. Scale bar: 50 µm. (C-D) Immunofluorescence localization of vimentin and E-cadherin (for fibroblasts and epithelial cells; C), and of F4/80 and E-cadherin (macrophages and epithelial cells; D) with a nuclear counterstain (Hoechst) in pancreatic sections from control, day 2, day 4, and day 6 mice. Scale bar: 50 µm. (F) Gene expression analysis by RT-qPCR of acinar (*Cpa1*, *Prss2*, *Amy2*) and ductal (*Krt19*, *Sox9*, *Onecut1*) markers, normalized to *18S* ribosomal RNA in pancreatic tissue from control, day 2, day 4, and day 6 mice. Data are shown as means ± SD. Statistical analysis: two-way ANOVA with Holm-Šídák correction. \**p* < 0.05, \*\**p* < 0.01, \*\*\**p* < 0.001, \*\*\*\**p* < 0.0001. n≥4.

**Supplementary Figure 2.**
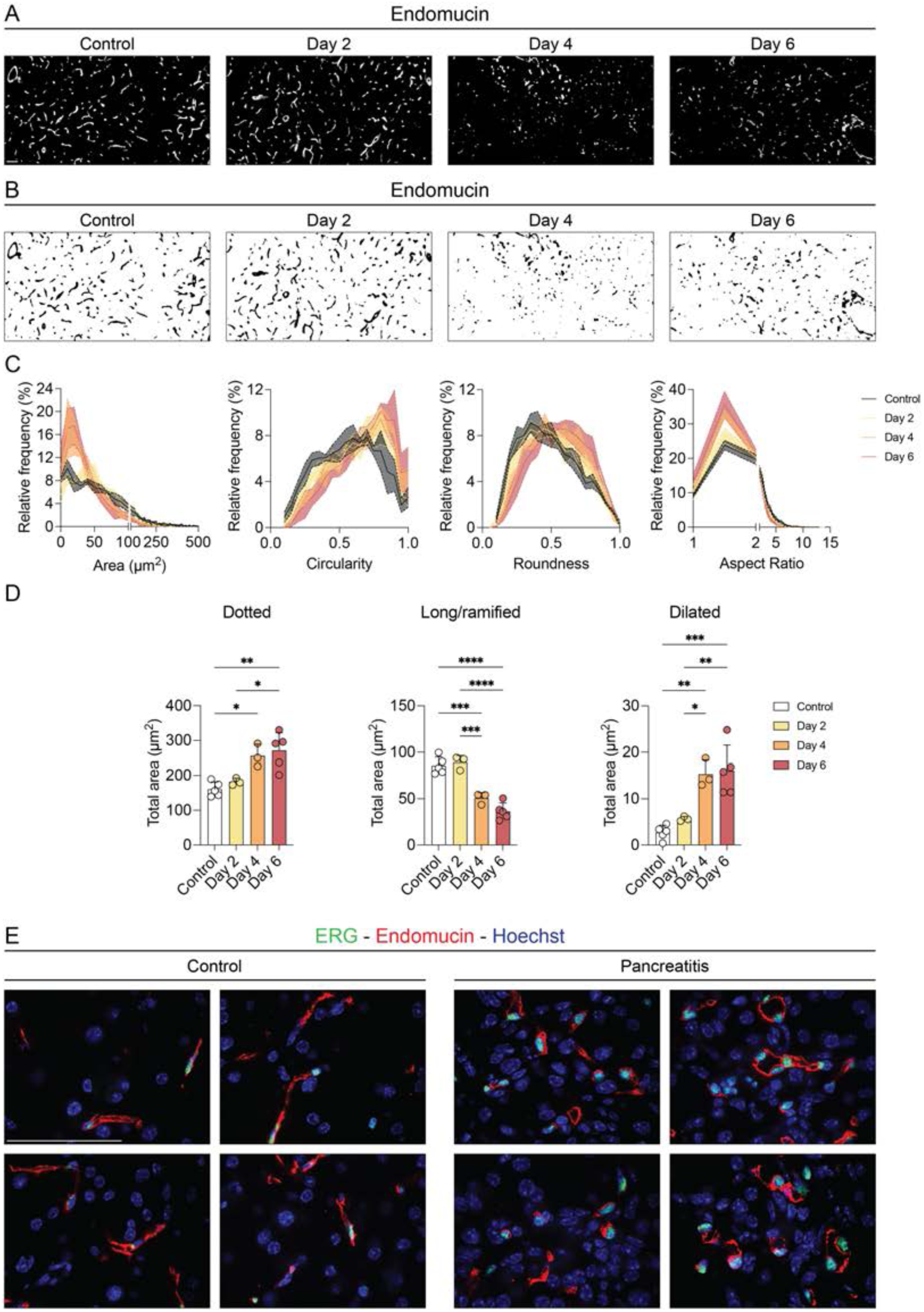
Blood vascular remodeling during the time-course of acute pancreatitis. (A) Immunofluorescence labeling of endomucin (endothelium) in pancreatic sections from control, day 2, day 4, and day 6 mice. Scale bar: 50 µm. (B) Segmentation of endomucin-positive objects from panel A for morphological analysis. (C) Quantification of the relative frequency (%) of endomucin-positive objects as a function of area, circularity, roundness, and aspect ratio in pancreatic sections from control, day 2, day 4, and day 6 mice. Data represent the cumulative frequency from five representative 488,160 µm^2^ fields per mouse. n=3 (day 2 and day 4), n=5 (control and day 6). (D) Quantification of endomucin-positive objects: dotted (area ≤ 80 µm^2^), long/ramified (area > 80 µm^2^, circularity < 0.7), and dilated (manually classified) in pancreatic sections from control, day 2, day 4, and day 6 mice. Data represent the average counts from five representative 488,160 µm^2^ fields per mouse and are shown as means ± SD. Statistical analysis: two-tailed Student’s t-test. \**p* < 0.05, \*\**p* < 0.01, \*\*\**p* < 0.001, \*\*\*\**p* < 0.0001. n=3 (day 2 and day 4), n=5 (control and day 6). (E) Immunofluorescence labeling of endomucin (endothelial cells) and ERG (endothelial cell nuclei), with a nuclear counterstain (Hoechst), and high-magnification visualization in pancreatic sections from control and caerulein-treated mice. Scale bar: 50 µm.

**Supplementary Figure 3.**
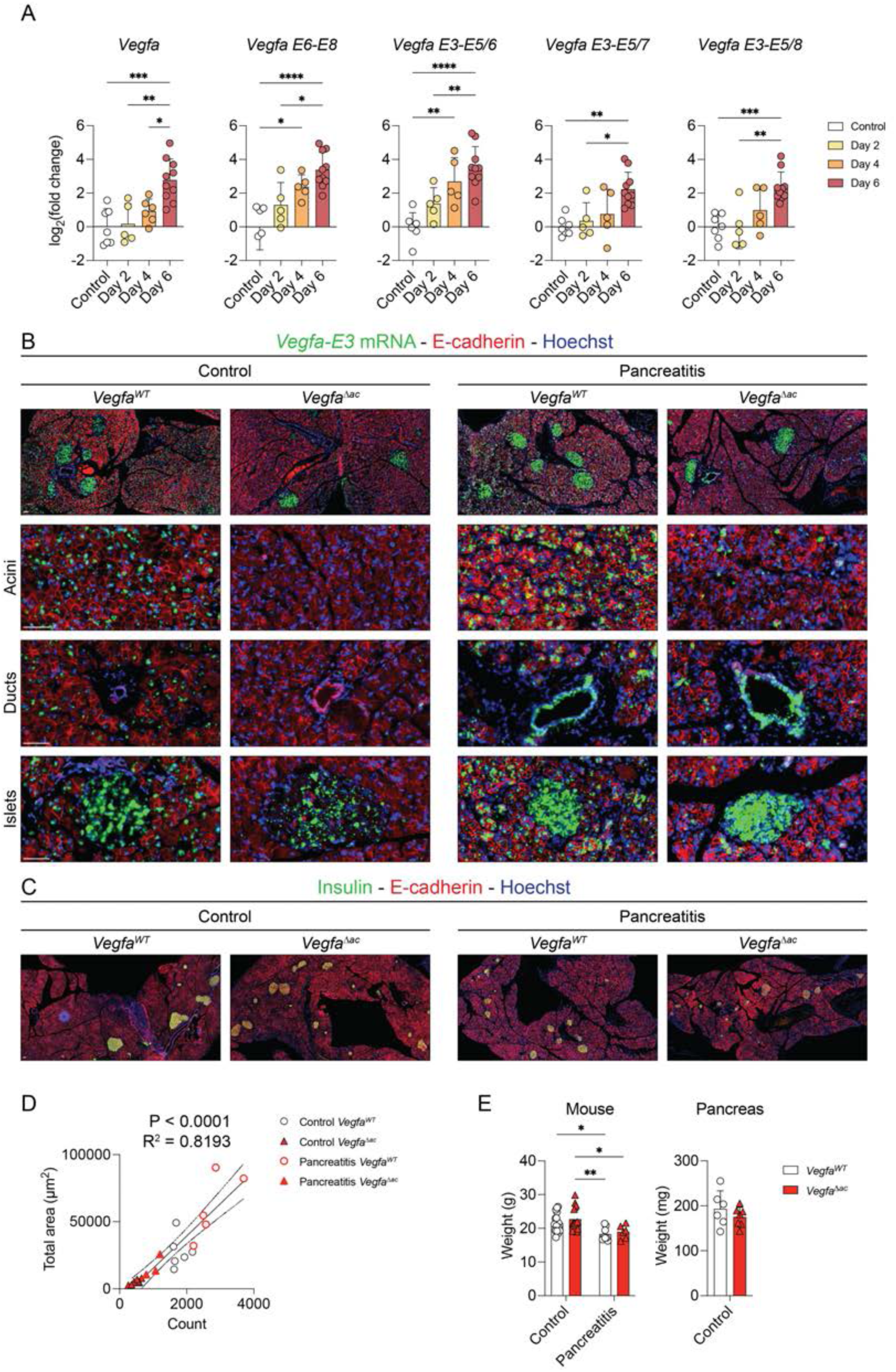
VEGF-A expression dynamics during the time-course of acute pancreatitis, spatial localization, and impacts on endocrine function and body weight. (A) Gene expression analysis by RT-qPCR of *Vegfa* and splice variants (E6-E8, E3-E5/E6, E3-E5/E7, E3-E5/E8), normalized to *18S* ribosomal RNA, in pancreatic tissue from control, day 2, day 4, and day 6 mice. Data are shown as means ± SD. Statistical analysis: two-tailed Student’s t-test. \**p* < 0.05, \*\**p* < 0.01, \*\*\**p* < 0.001, \*\*\*\**p* < 0.0001. n≥5. (B) *In situ* hybridization of *Vegfa-E3* by BaseScope™ and immunofluorescence labeling of E-cadherin (epithelial cells), with a nuclear counterstain (Hoechst) in pancreatic sections from control and caerulein-treated *Vegfa^WT^* and *Vegfa*^Δ*ac*^ mice. Close-ups display the acini, ducts, and islets of Langerhans. Scale bar: 50 µm. (C) Immunofluorescence labeling of Insulin (endocrine beta cells) and E-cadherin (epithelial cells), with a nuclear counterstain (Hoechst) in pancreatic sections from control and caerulein-treated mice. Scale bar: 50 µm. (D) Correlation between BaseScope™ *Vegfa-E3* object counting and total area, assessed using linear regression (straight middle line), with 95% confidence interval (curved lines around the middle line), Goodness of Fit (R^2^), and the significance of a non-zero slope (*p* value). (E) Body and pancreas weight from control and/or caerulein-treated *Vegfa^WT^* and *Vegfa*^Δ*ac*^ mice. Data are shown as means ± SD. Statistical analysis: two-way ANOVA with Holm-Šídák correction (mouse weight) or two-tailed Student’s t-test (pancreas weight). \**p* < 0.05, \*\**p* < 0.01, \*\*\**p* < 0.001, \*\*\*\**p* < 0.0001. n≥6.

**Supplementary Figure 4.**
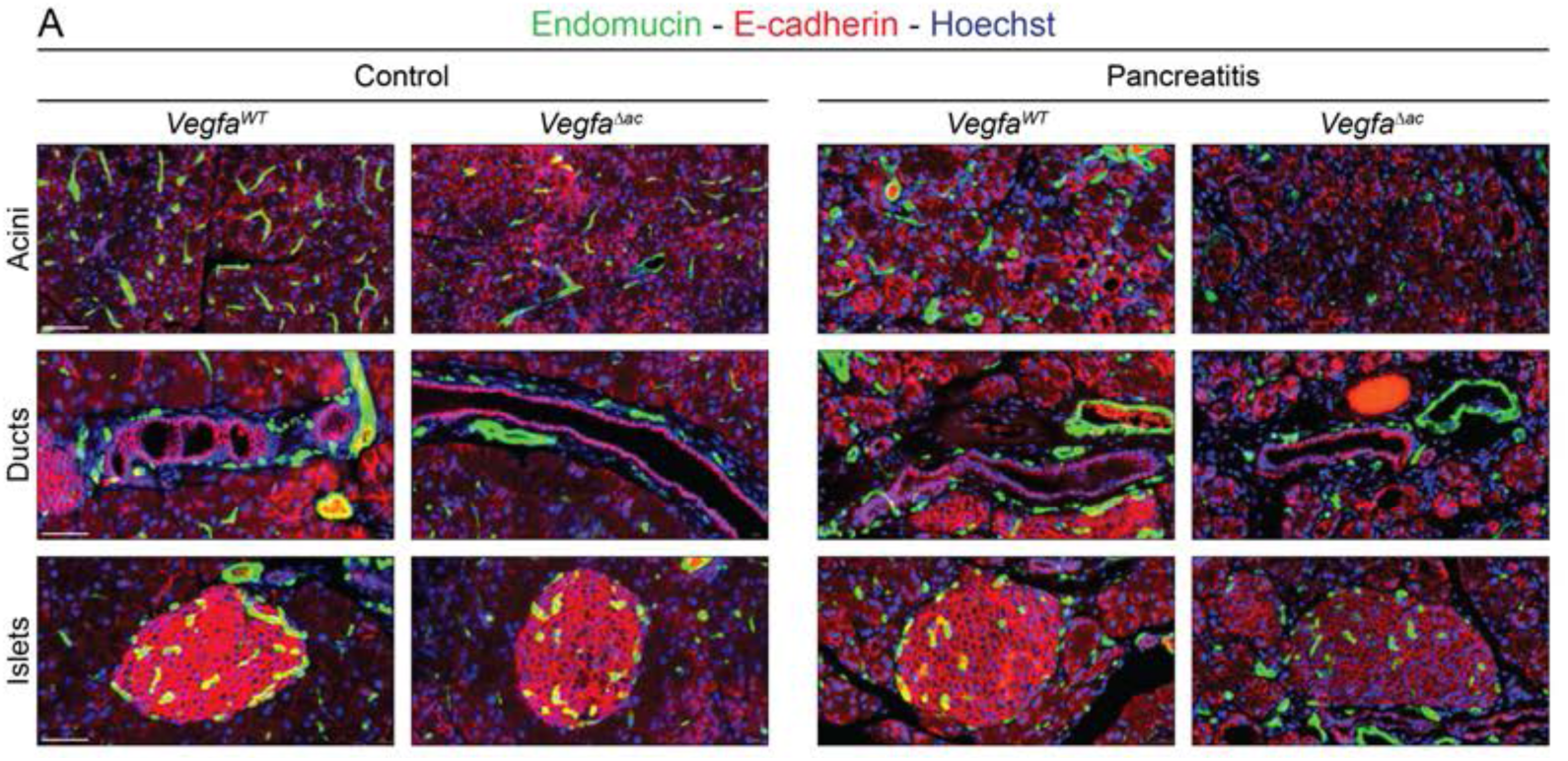
Blood vascular organization across pancreatic compartments and the impact of acinar-specific VEGF-A inactivation. Immunofluorescence labeling of endomucin (endothelial cells) and E-cadherin (epithelial cells), with a nuclear counterstain (Hoechst) in pancreatic sections from control and caerulein-treated *Vegfa^WT^* and *Vegfa*^Δ*ac*^ mice. Close-ups display the acini, ducts, and islets of Langerhans. Scale bar: 50 µm.

**Supplementary Figure 5.**
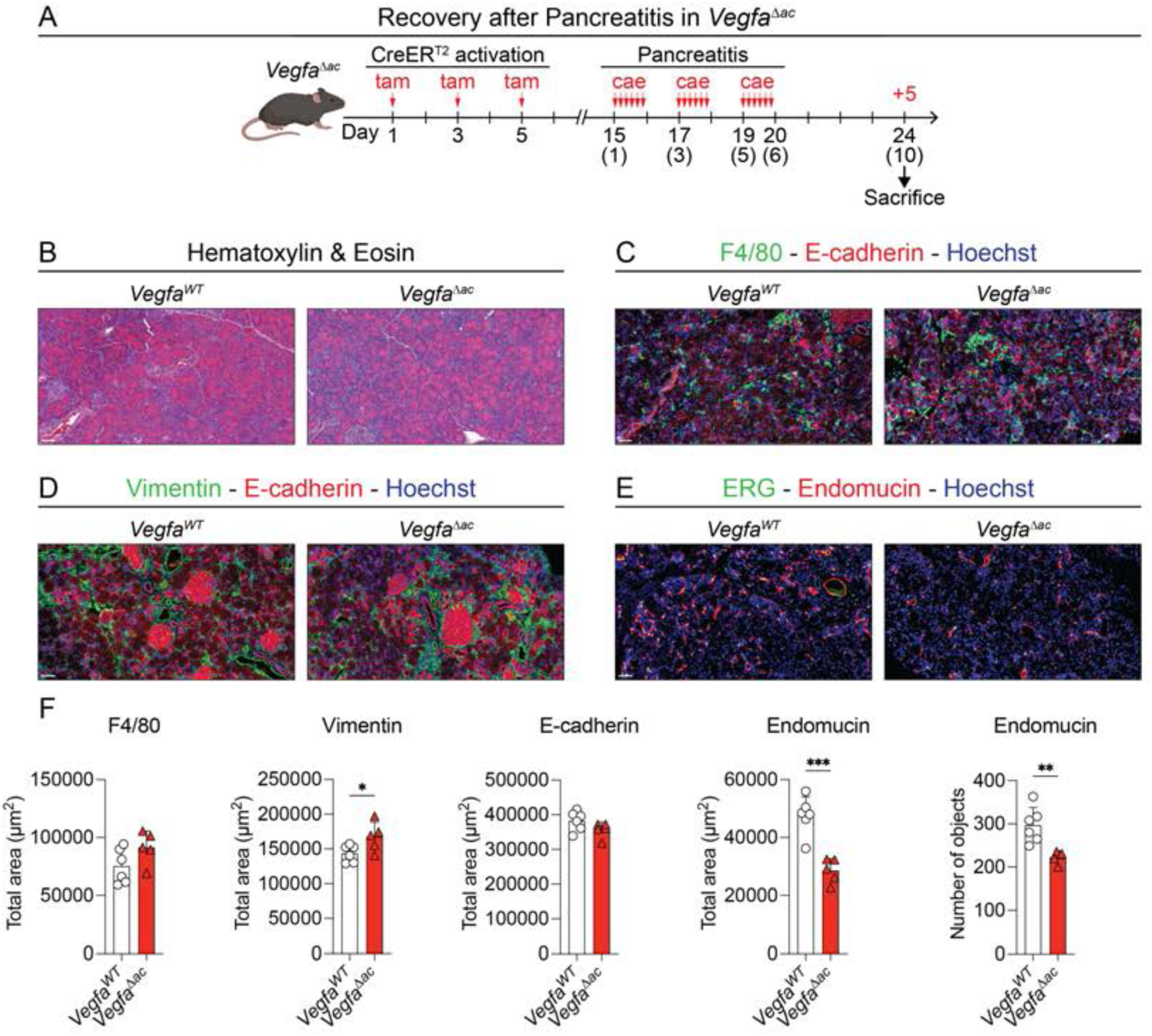
Effects of acinar-specific VEGF-A inactivation on regeneration post-acute pancreatitis. (A) Timeline of tamoxifen and caerulein injections (vertical red arrows), and sacrifice for tissue harvest on day +5 post-final caerulein injection (day 24 if counting from the initial tamoxifen injection) in *Vegfa^WT^* and *Vegfa*^Δ*ac*^ mice. (B) Hematoxylin (nuclei) & eosin (cytoplasm) staining of pancreatic sections from *Vegfa^WT^* and *Vegfa*^Δ*ac*^ mice at day +5 post-final caerulein injection. Scale bar: 50 µm. (C-E) Immunofluorescence labeling of F4/80 and E-cadherin (macrophages and epithelial cells; C), of vimentin and E-cadherin (for fibroblasts and epithelial cells; D), and of ERG and endomucin (for endothelial nuclei and cells); E), with a nuclear counterstain (Hoechst) in pancreatic sections from *Vegfa^WT^* and *Vegfa*^Δ*ac*^ mice at day +5 post-final caerulein injection. Scale bar: 50 µm. (F) Quantification of immunofluorescence labeling: total area (µm^2^) of F4/80, vimentin, E-cadherin, and endomucin, as well as the number of endomucin-positive objects, in pancreatic sections from *Vegfa^WT^* and *Vegfa*^Δ*ac*^ mice at day +5 post-final caerulein injection. Data represent the average total area from three representative 488,160 µm^2^ fields per mouse and are shown as means ± SD. Statistical analysis: two-tailed Student’s t-test. \**p* < 0.05, \*\**p* < 0.01, \*\*\**p* < 0.001, \*\*\*\**p* < 0.0001. n≥5.

## Notes

### Competing Interest Statement

The authors have declared no competing interest.

